# Paralogs of the *Candida albicans TLO* gene family form interconnected functional networks with incomplete redundancy

**DOI:** 10.64898/2026.06.29.735307

**Authors:** Emily Simonton, Nayeli Cangelosi, Mikki Zhou, Philip S. Hendricks, Andrew L. Woodruff, Matthew Z. Anderson

## Abstract

Gene duplication typically fails to confer a selective advantage to an organism, prompting their removal from a population. In the rare instance that duplication either does not incur a fitness cost or it enhances fitness, gene families can form through repeating the duplication process. While the function of gene duplicates has been studied in detail, little work has explored how repeated duplication impacts paralog redundancy and may restrict the emergence of new paralogs or novel function. Here, we constructed a panel of single deletion mutants for each of the 14 members of the *Candida albicans* te<u>lo</u>mere-associated (*TLO*) gene family to test the redundancy in molecular and biological function among paralogs from a lineage-specific expansion. Tlo proteins function as interchangeable subunits of the Mediator transcriptional regulatory complex and have the potential to alter gene expression and an array of cellular responses. Redundancy was the most common outcome, being observed for approximately 80% of the phenotypic assays in strains lacking single *TLO* genes. However, mutants for all 14 paralogs displayed non-redundant functions in phenotypes ranging from carbon utilization to *in vivo* virulence. Analysis of gene expression in single *TLO* mutants found similar trends in redundancy, and loss of single *TLO*s disproportionately affected genes involved in filamentation, adhesion, redox reactions, and transporter activity at the cell surface. Importantly, sequence divergence between paralogs positively correlated with the frequency of altered phenotypes in single *TLO* mutants, indicating the acquisition of non-redundant function with increased evolutionary distance. Double mutants lacking two *TLO* genes produced both positive and negative synergistic phenotypes, suggesting that crosstalk or coordinated regulation is common among paralogs. Together, this study demonstrates that recently emergent paralogs acquire non-redundant functions despite often retaining redundancy with other gene family members to form a highly interconnected functional network.

## INTRODUCTION

Most extant genes across the tree of life arose from other existing genes through a process of duplication and divergence. Gene duplication events predominantly result in loss or functional inactivation of the additional copy (Bergthorsson, et al. 2007; Adler, et al. 2014; Mihajlovic, et al. 2025). However, specialization for a prior minor function and/or acquisition of novel function through mutation, termed subfunctionalization and neofunctionalization, respectively, can favor paralog retention (Rastogi and Liberles 2005; Adler, et al. 2014; Andersson, et al. 2015). Eventually, these genes can be come fixed in a population through the selective advantages provided by these functions or genetic drift. Most investigations of paralog fates have been performed via the introduction of a second duplicate copy, disruption of a retained paralog following a whole genome duplication (WGD) event, or *in silico* surveys of duplicate outcomes after WGD (Kondrashov, et al. 2002; Dean, et al. 2008; Kassahn, et al. 2009; DeLuna, et al. 2010; Glasauer and Neuhauss 2014; Kuzmin, et al. 2020; Wilson and Liberles 2023; Mihajlovic, et al. 2025). While these studies explore gene fates when two copies are present, they do not account for the outcomes of repeated gene duplication that are critical to organism-specific adaptation and contribute to genome expansion.

Lineage-specific gene families composed of multiple paralogs can provide insight into the physiological processes under selection in an organism. Most of these gene families encode functionally constrained effectors that can be linked to organismal ecology and lifestyle (Mouches, et al. 1986; Romero and Palacios 1997; Rubin, et al. 2000; Wang, et al. 2019). For example, the *MAL*, *MEL*, and *SUC* genes in *Saccharomyces cerevisiae* can increase and decrease in copy number between strains depending on the availability of their carbon substrate (maltose, melibiose, and sucrose, respectively) (Codon, et al. 1998; Brown, et al. 2010; Wenger, et al. 2011). Gene families associated with organism-specific lifestyles in eukaryotes frequently reside in the subtelomeres, regions adjacent to the telomeric repeats that are rich in repetitive sequences and evolve quickly (Linardopoulou, et al. 2007; Brown, et al. 2010; Brann, et al. 2024). Recombination between subtelomeric repeats facilitates frequent gene loss and gain that can rapidly produce and remove paralogous genes (Linardopoulou, et al. 2005; Rudd, et al. 2007; Christiaens, et al. 2012; Anderson, et al. 2015; Freire-Beneitez, et al. 2016), while elevated mutation rates in the subtelomeres promote paralog diversification (Anderson, et al. 2008; Bergstrom, et al. 2014; Ene, et al. 2018). Consequently, subtelomeres are hotspots for the formation of large and genetically diverse gene families due to a combination of high rates of sequence evolution, genetic drift, and strong selective pressures.

There is a historical lack of systematic and focused dissection of gene families due to unique challenges when investigating large numbers of paralogs in a single organism. As the number of gene family members increase, there is an exponential increase in the number of mutants needed to isolate the function of each paralog through targeted deletions to capture all paralog interactions. This is further complicated by high sequence similarity between paralogs that can increase the likelihood of off-target effects when attempting to modify a specific paralog. Thus, most explorations of gene family paralog function focus on isolated representatives or assume that paralogs all contribute to a unified molecular or cellular function (Domergue, et al. 2005; Govender, et al. 2008; Wu, et al. 2013; Lemieux, et al. 2024). These approaches often fail to consider the functional interactions among paralogs and cannot measure the degree of redundancy among paralogous family members (Simon, et al. 2007; Li, et al. 2008; Correia, et al. 2010; Laruson, et al. 2020).

Expansion of the te<u>lo</u>mere-associated (*TLO*) gene family in the common fungal commensal and opportunistic pathogen *Candida albicans* provides an ideal platform to test redundancy among members of a lineage-specific gene family. The *TLO* gene family has experienced the greatest increase in copy number of any gene family in the most clinically relevant *Candida* species, *C. albicans* (Butler, et al. 2009; Schroeder, et al. 2025). Most *Candida* species encode a single *TLO* homolog, whereas the *C. albicans* genome contains 14 paralogs in the genome reference strain (Butler, et al. 2009; Jackson, et al. 2009). The *TLO* genes encode a MED2 domain that directs its incorporation into the major transcriptional regulatory complex, Mediator (Anderson, et al. 2012; Zhang, et al. 2012). Each Mediator complex contains a single copy of Tlo (or its ortholog Med2) (Zhang, et al. 2012; Uthe, et al. 2017), so that a Mediator molecule can only contain one Tlo paralog but different Mediator molecules in a cell can incorporate different Tlos. While the conserved MED2 domain defines the *TLO* paralogs as Mediator subunits, the genes cluster into three architectural groups (α, β, and γ) that are distinguished by differences in expression and gain and loss of genetic sequence in the 3’ end of the open reading frame (Anderson, et al. 2012; Dunn, et al. 2022). In general, *TLO*γ genes are expressed at levels 10-100x lower than either *TLO*α or *TLO*β genes at the transcript and protein level across multiple conditions (Anderson, et al. 2012; Zhang, et al. 2012; Anderson, et al. 2014). Within an architectural group, *TLO*s contain ∼97% nucleotide identity, and 82% identity exists between architectural groups, excluding indels (Anderson, et al. 2012). All *TLO* genes reside in the subtelomeres of the *C. albicans* chromosomes apart from a single chromosomal internal copy, *TLO*α*34*. *TLO* expansion is likely a consequence of frequent recombination among *C. albicans* subtelomeres that continues to contribute to copy number variation seen among clinical isolates (Anderson, et al. 2015; Hirakawa, et al. 2015; Dunn, et al. 2022).

Construction of a *tlo*Δ/Δ mutant (lacking all 14 *TLO* genes) in *C. albicans* revealed pleiotropic transcriptional and phenotypic roles for the gene family (Fletcher, et al. 2023). In contrast, loss of only *TLO*β*2* resulted in more limited phenotypic consequences related to filamentation and biofilm formation defects (Uppuluri, et al. 2018), indicative of some degree of functional redundancy with other *TLO* paralogs. Here, we systematically explored functional redundancy among all members of the *TLO* gene family by constructing a panel of mutants lacking one of the 14 paralogs. Mutants for single *TLO* genes lacked evidence of altered phenotypes in most conditions but displayed an increased prevalence of mutant phenotypes under stress. The frequency of phenotypic changes caused by loss of a single paralog was positively correlated with the number of new sequence variants. Yet, single *TLO* mutants lacking any discernable cellular phenotype under a specific condition had altered expression of 10s to 100s of genes compared to the wildtype strain. Furthermore, disruption of two *TLO* genes unveiled complex epistatic interactions among paralogs beyond redundancy. Taken together, this study demonstrates that individual *TLO* paralogs retained substantial functional overlap during expansion but also acquired unique molecular and physiological roles to produce an interwoven meshwork of paralog-phenotype interactions, where multiple paralogs impact a single trait and single paralogs exhibit extensive pleiotropy.

## RESULTS

A previous investigation constructed a complete *tlo* null mutant (*tlo*Δ/Δ) by targeting the conserved MED2 domain of *TLO* genes for disruption with CRISPR/Cas9. However, *tlo*Δ/Δ mutants contained unintentional chromosomal truncations, fusions, and translocations that may significantly impact phenotypic expression (Fletcher, et al. 2023). All *tlo*Δ/Δ mutants had chromosomal rearrangements between long terminal repeat (LTR) elements at the chromosomal-internal *TLO*α*34* locus on chromosome 1 (Chr1) and in the subtelomeres. Therefore, we rebuilt a *tlo*Δ/Δ strain lacking any significant karyotypic aberrations by first deleting the *TLO*α*34* locus with CRISPR/Cas9 (Fig 1A). Following construction of two *tlo*α*34*Δ/Δ mutants, we deleted the remaining subtelomeric *TLO* loci with CRISPR/Cas9 disruption of the conserved N-terminal MED2 domain. Loss of all *TLO* genes was confirmed by PCR and whole genome sequencing. The two independent *tlo*Δ/Δ lineages lacked any significant karyotypic changes with only a few instances of subtelomeric exchanges that were telomere-proximal to any known protein-coding genes.

**Fig. 1:**
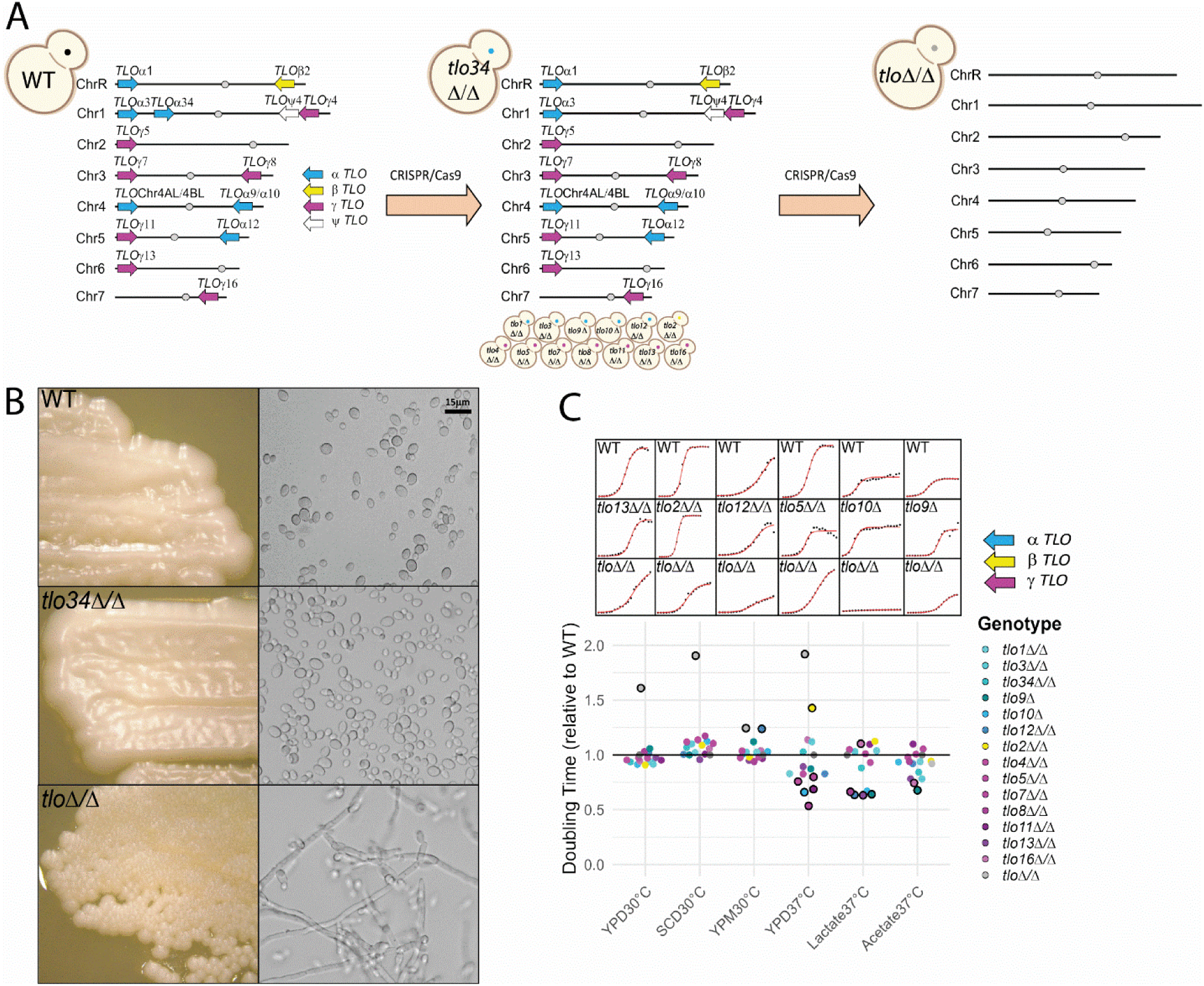
*TLO* genes display non-redundant roles in sub-optimal carbon sources. (**A**) Mutants lacking single *TLO* genes were constructed via CRISPR/Cas9-mediated targeted deletions. The *tlo* null (*tlo*Δ/Δ) was built from the *tlo*α*34*Δ/Δ strains by targeting a conserved region of the remaining *TLO* paralogs. (**B**) The indicated strains were grown on solid YPD medium at 30°C for 2 days, and images of the colony and cellular morphologies were captured at 5x and 40x magnification, respectively. Scale bar = 15 μm. (**C**) Doubling times for the wildtype, single *TLO* mutants, and the *tlo*Δ/Δ strain grown for 18-72 hours (until growth curves reached maximum OD and plateaued) in liquid yeast peptone medium supplemented with 2% of the indicated carbon source, apart from lactate at 0.4% and normalized to the mean WT growth. Representative growth curves are shown above each condition. *TLO* architectural groups are color-coded as indicated, and the *tlo*Δ/Δ strain is represented by light gray. Significance determined by a modified Student’s t-test at p≤0.05 is indicated by a bold outline. N≥4. **Fig. 1 alt text:** Comparison of single *TLO* mutants versus the complete knockout strain via karyotype graphical representation, colony and cellular morphology, and normalized doubling times in six different growth conditions.

Assaying a *tlo*Δ/Δ mutant allows for broad definitions of *TLO* function but does not provide insight into functional redundancy among individual paralogs. To assess *TLO* redundancy, we produced a panel of single *TLO* deletion mutants using CRISPR/Cas9. Sequence conservation among the *TLO* coding sequences is too high to confidently target a single open reading frame (ORF), so we designed guide RNAs (gRNAs) to cleave sequence-specific regions immediately flanking the *TLO* gene of interest. Repair templates spanned the cleavage site and the target *TLO* locus, producing strains containing the precise desired gene deletion with no detectable karyotypic variation (Fig 1A, S1). We applied this approach to all 14 highly homologous *TLO*s in SC5314 to obtain a panel of single *TLO* deletion mutants with two independently obtained mutants per *TLO* gene.

The previously constructed *tlo*Δ/Δ mutants grew primarily as wrinkled colonies composed of pseudohyphae on YPD plates at 30°C (Fletcher, et al. 2023). Both of the newly produced *tlo*Δ/Δ strains lacking chromosomal rearrangements also grew as wrinkled colonies composed primarily of pseudohyphal cells (Fig 1B). In contrast, strains lacking only one *TLO* gene formed smooth colonies composed of yeast cells, similar to the wildtype strain. We can conclude that the altered morphology of the *tlo* null mutant is not dependent on any single paralog.

### *TLO* genes display non-redundant functions for growth in organic acids

In other Ascomycetes, carbon sources can select for copy number variation in subtelomeric expanded gene families that contribute to nutrient uptake and metabolism, which promotes growth (Winzeler, et al. 2003; Brown, et al. 2010). Growth of the *tlo*Δ/Δ mutant was significantly reduced in YPD and SCD rich media at 30°C compared to the wildtype SC5314 parental strain (WT), suggesting that *TLO*s also contribute to metabolic regulation (Fig 1C, S2). However, loss of any single *TLO* gene had a negligible effect on growth rates under the same rich media conditions. Replacement of dextrose with the monosaccharide fructose or the disaccharide sucrose produced comparable results: The *tlo*Δ/Δ mutant displayed a slow growth phenotype and strains lacking single *TLO* genes grew similar to the WT (Fig S2). Growth in medium with maltose as the main carbon source (YPM) increased the doubling time of the WT strain two-fold. A growth defect remained for the *tlo*Δ/Δ mutant in YPM but was substantially more similar to the WT (Fig 1C). Importantly, loss of *TLO*α*12* alone also produced a growth defect indistinguishable from the *tlo*Δ/Δ strain (Student’s t-test, *t*(62)=2.89, p=0.005), whereas no other single *TLO* mutant impacted growth under these conditions. This suggests widespread redundancy among *TLO* genes for use of most carbon sources, and the potential for non-redundant functions to emerge for single paralogs.

Loss of single *TLO* genes had a greater impact when grown on non-fermentable carbon sources. Consistent with the literature (Priest and Lorenz 2015), the growth rate of WT cells was dramatically reduced in medium containing acetate or lactate as the primary carbon source relative to media containing sugars (Fig S2). The *tlo*Δ/Δ mutant grew as well as the WT in acetate but failed to produce any detectable growth in lactate (Fig 1C). Only two mutants (*tlo*α*9*Δ and *tlo*γ*16*Δ/Δ) displayed an acetate-related growth phenotype, but loss of six separate individual *TLO*s altered growth in lactate. In acetate, the two *TLO* mutants displayed decreased doubling times that were not observed in the *tlo*Δ/Δ mutant, suggesting that loss of additional paralogs negates these minor growth advantages. In contrast, loss of *TLO* genes both increased and decreased growth rates in lactate medium, and, surprisingly, most mutants displayed increasing growth rates as observed in acetate. However, the lack of noticeable growth of the *tlo*Δ/Δ mutant suggests that *TLOs* are indispensable for lactate utilization. Incubation of *TLO* mutants in artificial saliva medium displayed the opposite effects to lactate. Nine of 14 single *TLO* mutants showed attenuated growth in saliva medium, but the *tlo*Δ/Δ mutant had increased growth rates (Fig S2). These data suggest a complex interplay among *TLO* genes in the regulation of non-fermentable carbon metabolism that can produce bidirectional phenotypic changes based on loss of single genes or all paralogs together.

Increasing the temperature to 37°C during growth in liquid YPD medium had a substantial impact on cell behavior of single *TLO* mutants. Loss of *TLO*γ*5 TLO*γ*7*, *TLO*γ*8*, *TLO*α*10*, or *TLO*γ*11* increased the growth rate of *C. albicans* compared to the WT, whereas the *tlo*β*2*Δ/Δ and *tlo*Δ/Δ mutants displayed decreased growth rates (Fig 1C, S2). Taken together, large bidirectional changes to growth among single *TLO* gene mutants in this condition strengthened the evidence for complex regulatory relationships among *TLO* paralogs and their target loci under a single condition.

### *TLO*s play redundant functions under stress except phosphate starvation

*C. albicans* is exposed to a variety of stresses during colonization of the human body that can include high temperature, cell wall damaging agents, osmotic regulators, reactive oxygen species, and antifungal drugs (Brown, et al. 2014). Loss of all *TLO* genes led to high sensitivity towards most stressors (*e.g.*, Calcofluor White (CFW), heat shock at 42°C, tert-Butyl hydroperoxide (tBOOH), propiconazole) except for 1M sodium chloride (Fig 2, S3). For example, the *tlo*Δ/Δ mutant failed to grow when incubated with 25 μg/mL CFW in YPD at 30°C. Consistent with the previous description of increased resistance to fluconazole in a *tlo* null mutant (O’Connor-Moneley, et al. 2024), the *tlo*Δ/Δ strain displayed elevated resistance to propiconazole (Fig S3).

**Fig. 2:**
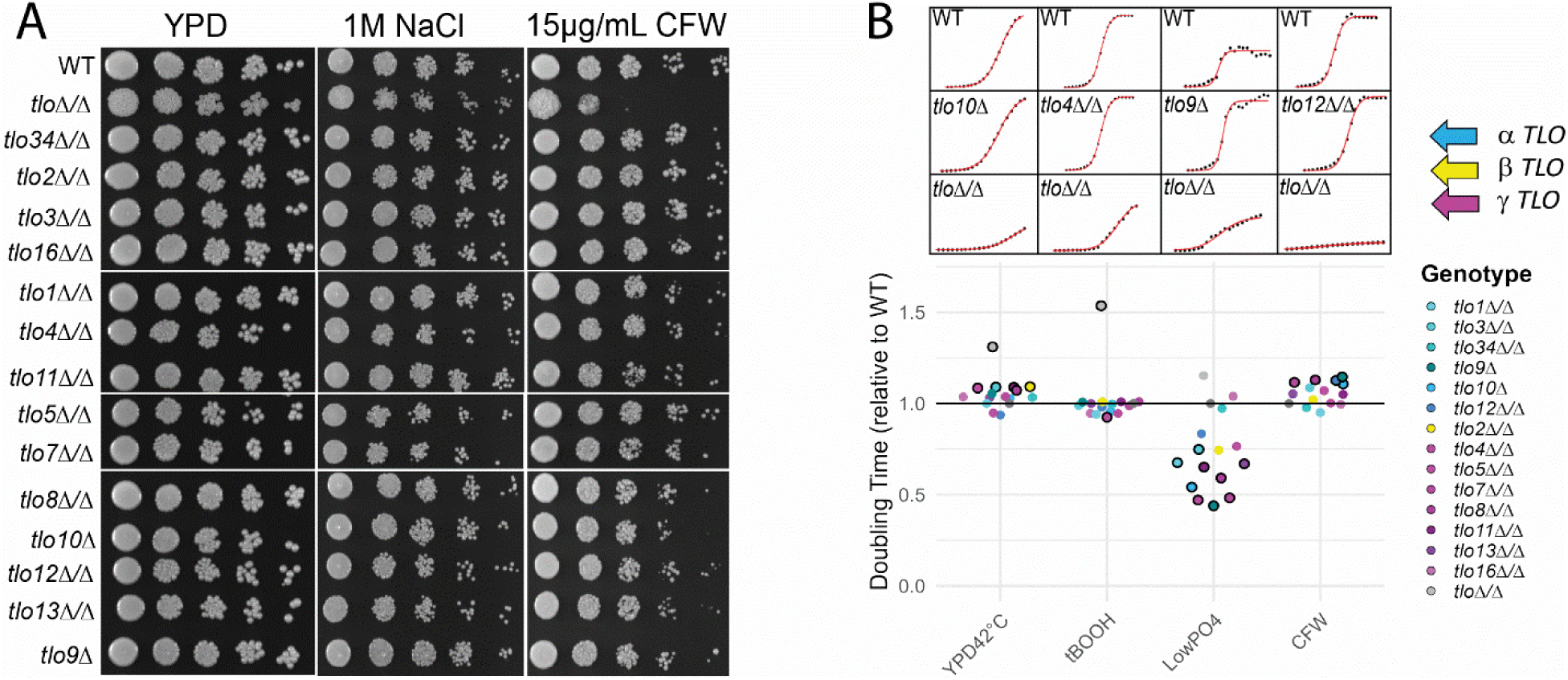
Stress resistance is largely redundant among *TLO* genes apart from low phosphate. (**A**) Strains were grown overnight, counted, and diluted to 1×10^7^ cells/mL. Cells were spotted as 3 μL aliquots in dilutions of 1:10 for the second dilution and then 1:5 from left to right on the indicated medium. Cells were incubated at 30°C for 48 hours prior to imaging (**B**) Doubling times for strains grown for 24 hours at 30°C in liquid YPD+0.3 mM tert-butyl hydroperoxide (tBOOH), in YPD+25 μg/mL CFW, in SCD+0.05 mM PO_4_ as well as liquid YPD at 42°C and normalized to the mean WT growth. Representative growth curves are shown above each condition. *TLO* architectural groups are color-coded as indicated, and the *tlo*Δ/Δ strain is represented by light gray. Significance by a modified Student’s t-test at p≤0.05 is indicated by a bold outline. N≥3. **Fig. 2 alt text:** Spot dilution of *TLO* mutant cells grown in osmotic and cell wall stress conditions are shown alongside normalized doubling time data in four different stress conditions, with subfigures labelled a and b.

In contrast to the *tlo*Δ/Δ strain, single *TLO* mutants had few and relatively minor stress susceptibilities. Loss of a few *TLO* genes produced minor growth defects at 42°C or in the presence of CFW at 30°C in rich medium. Only the *tlo*γ*8*Δ/Δ strain had growth defects under more than one condition in rich medium (42°C and CFW), and the *tlo*γ*4*Δ/Δ strain had increased growth rates in tBOOH and decreased growth rates at 42°C. However, low phosphate conditions (0.05 mM v. 7.4 mM in SCD) nearly doubled the growth rates of multiple single *TLO* mutants compared to the WT strain (Fig S3). The relatively faster growth rates of the single *TLO* mutants were due to an increased doubling time of the WT parental strain in low phosphate conditions, which then mirrored the growth rate of the *tlo*Δ/Δ strain. *TLO*s were confirmed as drivers of observed phenotypes via single gene complementation in key mutant strains (Fig. S4). These results suggest that multiple *TLO*s may negatively regulate processes that use phosphate for growth independently or engage in crosstalk to regulate phosphate through a select subset of *TLO* gene(s).

### Single *TLO* mutants display defects in filamentation processes

Hyphal formation can be triggered by a diverse set of environmental stimuli that act upon interconnected signaling cascades (Noble, et al. 2017). This led us to hypothesize that loss of single *TLO* genes would show complex interactions as observed for growth in different media conditions. To test this hypothesis, yeast cells from the wildtype, *tlo*Δ/Δ, and the panel of single *TLO* mutants were incubated in liquid RPMI 1640 medium supplemented with 10% fetal bovine serum (FBS) or plated to solid Spider medium at 37°C. Consistent with previous reports (Fletcher, et al. 2023), approximately 50% of *tlo*Δ/Δ cells failed to form hyphae in liquid RPMI+10% FBS (Fig 3A, B). Only the *tlo*γ*5*Δ/Δ mutant displayed reduced filamentation compared to the wildtype parental strain under these conditions, suggesting it contains a distinct function in hyphal regulation compared to other *TLO* paralogs.

**Fig. 3:**
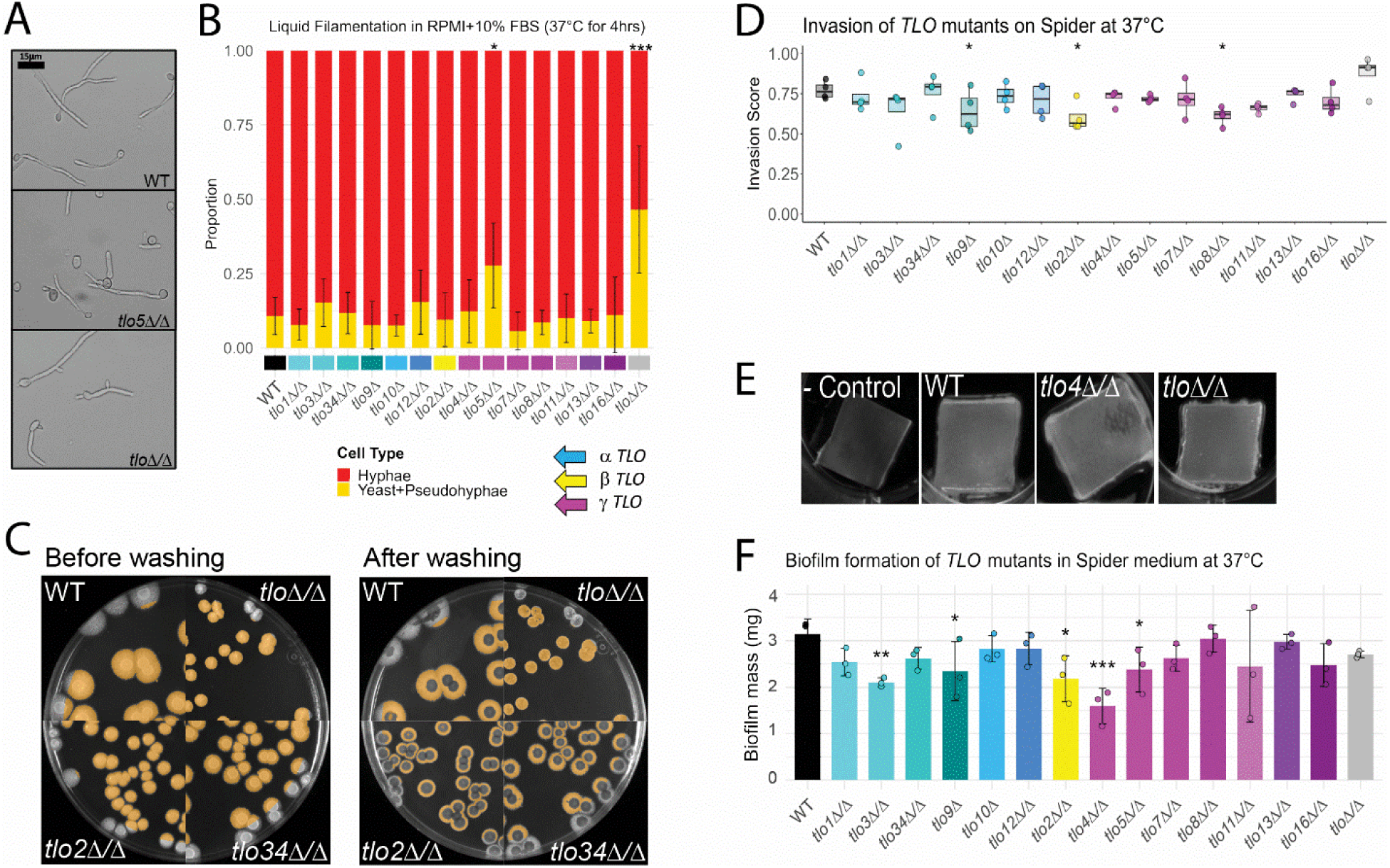
Filamentation and biofilm phenotypes do not follow *TLO* architectural groups. (**A**) Cells were grown in liquid RPMI 1640 + 10% fetal bovine serum (FBS) at 37°C for 4 hours and imaged at 40x magnification. Scale bar = 15 μm. (**B**) Cells were categorized as either true hyphae or yeast/pseudohyphae across at least 4 full fields of view with a minimum of 50 cells total. Data are plotted as the mean and standard deviation. N=4. (**C**) One hundred cells were plated to solid Spider plates from overnight cultures and allowed to grow as colonies at 37°C for 7 days before imaging. Plate images show colonies before and after washing in a stream of water for 3-5 seconds. Detected hyphae are highlighted in orange for colonies that do not contact the edge of the plate. (**D**) Quantification of the invasive fraction of the colony area for imaged plates. Boxplots are shown as the average and the interquartile ranges (IQRs) with whiskers extending to 1.5*IQR. N=4. (**E**) One half of an OD_600_ of cells were incubated with silicon elastomer squares pre-treated with adult bovine serum (ABS) and incubated in Spider medium for 90 min. Silicon squares were washed in 2 mL 1xPBS to remove non-adherent cells before incubating in fresh Spider medium for 66 hours at 37°C and imaged. (**F**) The dry weight of the biofilm formed on the silicone squares is plotted as the average and standard deviation. N=3. Significance was calculated via a modified Student’s t-test; * p < 0.05, ** p < 0.01, *** p < 0.001. **Fig. 3 alt text:** Filamentation-related data across subfigures labelled a to f, which demonstrate proportions of hyphal formation, adhesion and invasion on solid medium, and biofilm mass formation differences across single *TLO* mutants.

Plate-based assays provide the ability to assess colony adhesion, invasion, and filamentation, which can differ between solid and liquid forms (Azadmanesh, et al. 2017; Dunn, et al. 2020). Contrary to liquid filamentation, filamentation of *tlo*Δ/Δ colonies resembled the wildtype strain on solid Spider medium (Fig 3C, S5). Neither the *tlo*Δ/Δ mutant nor single *TLO* mutants displayed defects in colony adhesion or radial filamentation on Spider medium, but mutants from all three *TLO* architectural groups (*tlo*α*9*Δ/Δ, *tlo*β*2*Δ/Δ, and *tlo*γ*8*Δ/Δ) displayed reduced invasion of the agar substrate, indicative of non-redundant roles in a specific phase of invasive growth.

*C. albicans* forms highly structured biofilms through the collective actions of adhesion, invasion, and filamentation (Dunn, et al. 2020). To assess the contribution of *TLO* genes to biofilm formation, square punches of silicone elastomer used to make indwelling catheters were incubated with cells from the wildtype strain or each of the *TLO* mutants at 37°C in liquid Spider medium. After 66 hours (hrs) of incubation, the biofilm of the *tlo*Δ/Δ strain was similar in mass to the WT, whereas loss of *TLO*α*3, TLO*β*2, TLO*γ*4,* or *TLO*γ*5* reduced the biofilm mass by up to 50% (Fig 3E, F). Reduced biofilm formation of *tlo*γ*5*Δ/Δ and *tlo*β*2*Δ/Δ could be attributed to reduced filamentation and invasion, respectively. However, defects in biofilm formation of *tlo*α*3*Δ/Δ and *tlo*γ*4*Δ/Δ indicate a context-dependent role for these phenotypes that is reliant on the induction stimulus.

### Loss of single *TLO* genes increases virulence

Macrophages and other professional phagocytes engulf and destroy *C. albicans* cells during invasive overgrowth to restrict fungal dissemination and systemic disease. To determine the degree of redundancy among *TLO* genes for direct host-fungal interactions, immortalized RAW 264.7 mouse macrophages were incubated with *C. albicans* cells and allowed to interact for 20 hours before collecting supernatants to measure lactate dehydrogenase (LDH) release as a proxy for macrophage cell death (Fig 4A). The *tlo*Δ/Δ mutant caused significantly less LDH release from RAW264.7 cells, reflecting a defect in pathogenesis (Fig 4B). In contrast, loss of five individual *TLO* genes (*TLO*α*9*, *TLO*γ*5*, *TLO*γ*8*, *TLO*γ*11*, or *TLO*γ*13*) approximately doubled the release of LDH from RAW 264.7 cells, indicative of elevated *TLO* mutant pathogenesis.

**Fig. 4:**
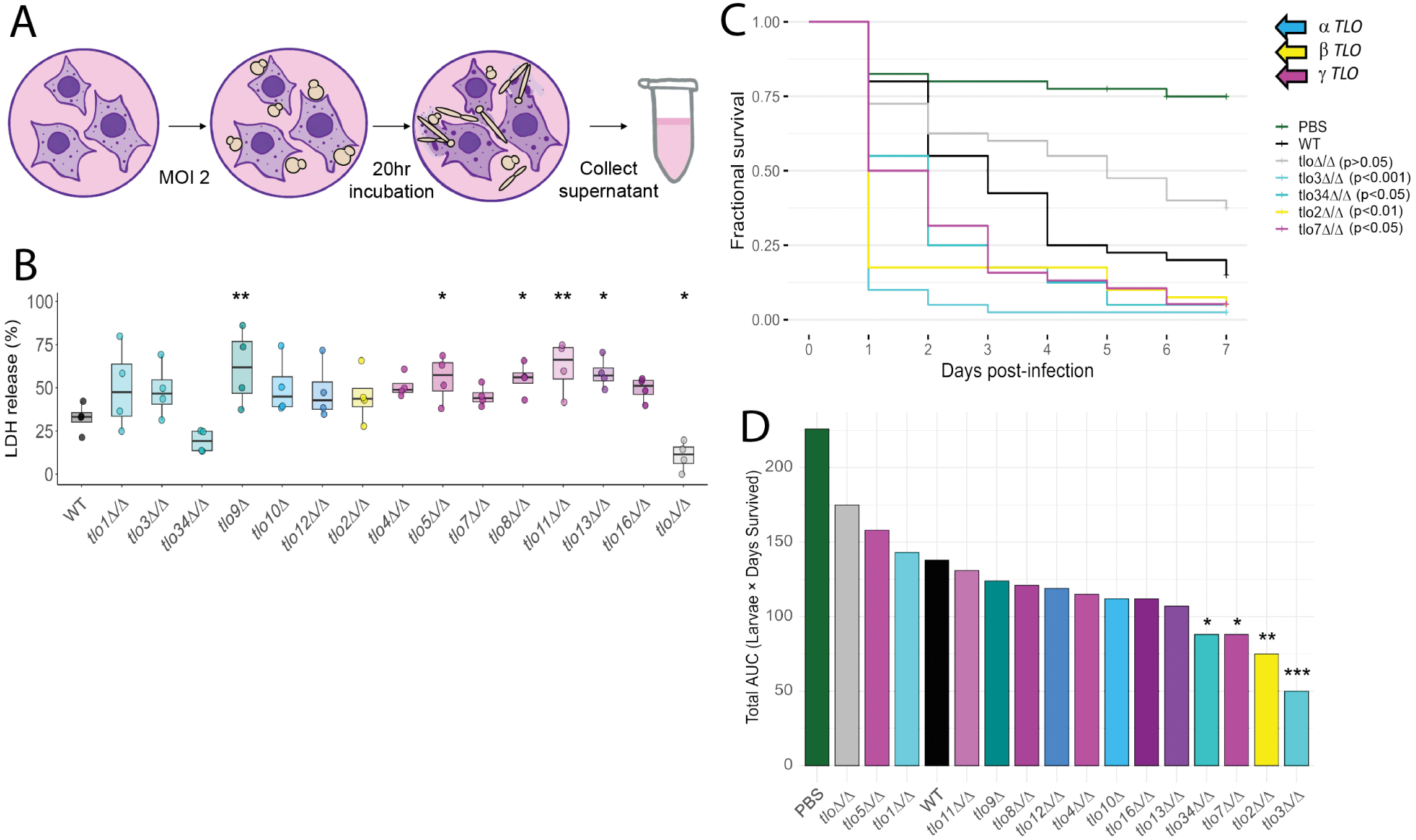
Loss of single *TLO* genes increases virulence phenotypes. (**A**) RAW 264.7 murine macrophages were incubated with *C. albicans* cells at an MOI of 2. After 20 hours, supernatants were collected, and lactate dehydrogenase (LDH) release was measured. (**B**) Calculated LDH release for the indicated strains normalized to negative and positive cell lysis controls as the minimum and maximum LDH release. N=4. Significance was determined via modified Student’s t-tests; * p < 0.05, ** p < 0.01, *** p < 0.001. (**C**) *Galleria mellonella* survival is plotted following infections with 2.5×10^5^ cells for the indicated *C. albicans* strain or PBS as a negative control. (**D**) The area under the curve (AUC) was calculated for the combined Kaplan Meier plots of *G. mellonella* infection with *C. albicans*. N=40 animals. Significance was determined via a Log-Rank test with p-value corrections performed by Benjamini-Hochberg method; * p < 0.05, ** p < 0.01, *** p < 0.001. **Fig. 4 alt text:** Graphical methods summary of infection and LDH release by murine macrophages inoculated with single *TLO* mutants to test *in vitro* virulence. *In vivo* virulence was demonstrated by infection of wax worm larvae and monitoring survival across seven days, with subfigures labelled a to d.

To identify changes in virulence due to loss of single *TLO* genes, the wildtype, *tlo*Δ/Δ mutant, and single *TLO* deletion mutants were individually injected into the waxworm *Galleria mellonella* to model disseminated candidiasis (Hirakawa, et al. 2015; Dunn, Woodruff, et al. 2018). As in the macrophage killing assay, the *tlo*Δ/Δ strain was less virulent in infected larvae compared to the wildtype background strain. However, a different set of *TLO* genes displayed altered virulence in this model of systemic disease compared to macrophage killing. Injection of the *tlo*α*3*Δ/Δ, *tlo*α*34*Δ/Δ*, tlo*β*2*Δ/Δ, or *tlo*γ*7*Δ/Δ mutants increased virulence compared to injection of the wildtype (Fig 4C, D). Together, this data suggests that *TLO*s play non-redundant roles in suppressing pathogenesis in the host but are collectively important for causing disease in the host.

### Non-redundant *TLO* functions are associated with sequence evolution but do not cluster by architectural group

To interpret the phenotypic profiles across *TLO* mutants, we built a heatmap for all assayed phenotypes that was color-coded by significant differences relative to WT. The *tlo*Δ/Δ mutant performed poorly under most (14/22) assayed conditions, indicating a general fitness defect that was not present for mutants of single *TLO* genes. Loss of single *TLO* genes failed to produce statistically different phenotypes in most cases (246/308 potential phenotype x strain interactions), indicating widespread redundancy (Fig 5A). Yet, each mutant for a single *TLO* gene displayed between one and eight distinct phenotypes relative to WT. Mutants for two *TLO*α paralogs, *TLO*α*1* and *TLO*α*34*, generally behaved similarly to the wildtype with only one phenotypic change, whereas *tlo*α*9*Δ and *tlo*γ*8*Δ/Δ strains exhibited alterations to eight phenotypes compared to the wildtype strain. Phenotypes produced by the loss of single *TLO*s generally clustered around growth based on nutrient availability, filamentation, and pathogenesis. Unidirectional phenotypic changes were generally observed for *TLO* mutants that were exposed to stressors, induced to form biofilms, or assayed for pathogenesis, whereas growth rates of *TLO* mutants changed bidirectionally between and within a single condition.

**Fig. 5:**
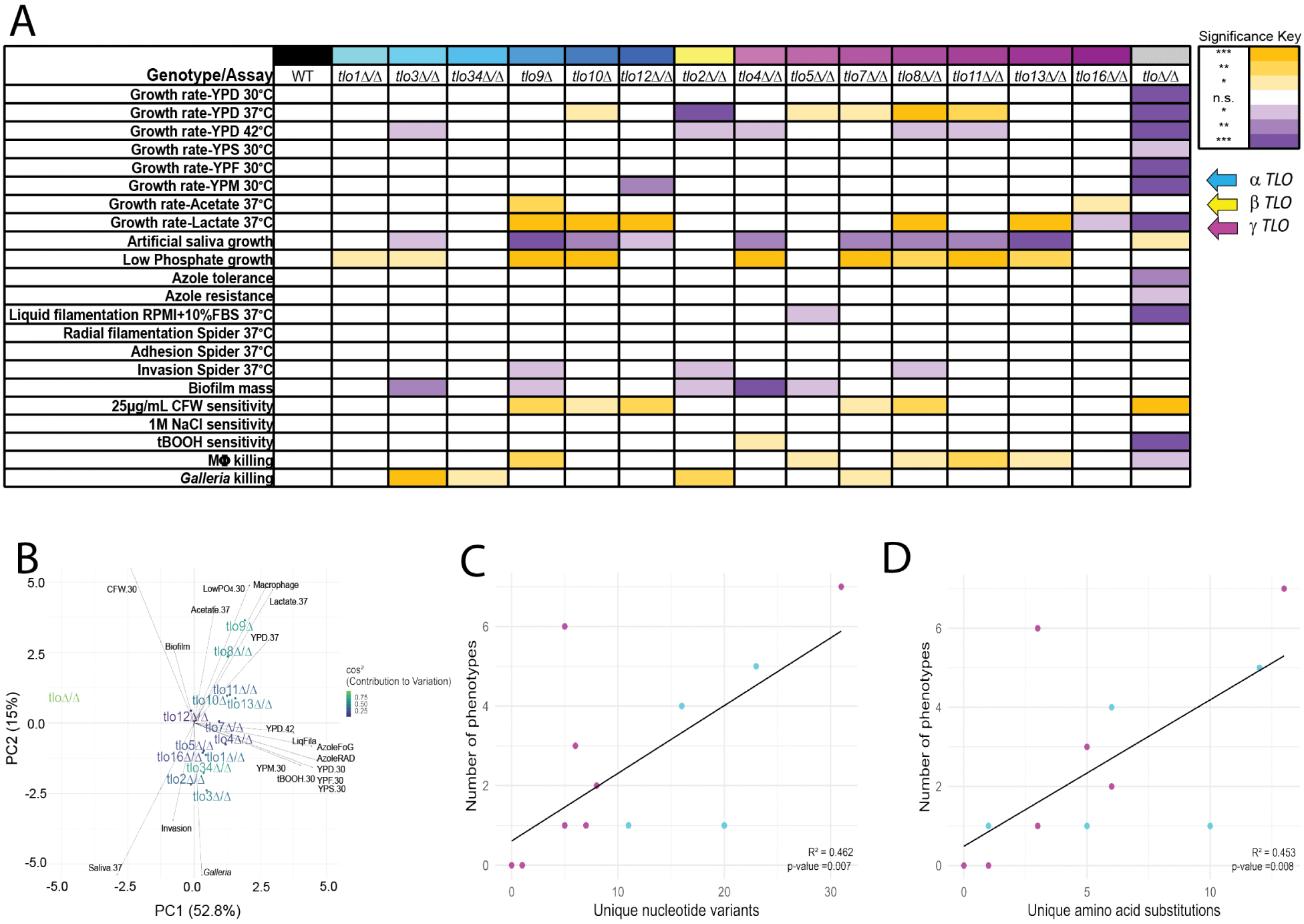
The frequency of *TLO* mutant phenotypes correlate with genetic distance but not the architectural group. (**A**) All single *TLO* mutants and the *tlo*Δ/Δ are shown with their relative phenotypic differences compared to WT. White indicates no significant difference from WT, whereas increasing or decreasing phenotypes are indicated by increasing shades of orange or purple, respectively. (**B**) The normalized significance for each genotype across phenotypic assays was used to construct a PCA biplot. Vectors indicate the magnitude and direction of each assay. *TLO* mutants are plotted to indicate their contributions to the variance and color-coded to indicate their relative contribution. (**C, D**) A phylogenetic reconstruction of *TLO* relatedness was built for each *TLO* architectural group using a distance-matrix UPGMA (Unweighted Pair Group Method using arithmetic Averages). The number of unique nucleotide variants and amino acid substitutions that segregated exactly at each branchpoint were counted. The number of sequence changes was plotted against the phenotypes that segregate at the same branchpoint. **Fig. 5 alt text:** Graphics and data in subfigures are labelled a to d and summarize significant phenotypes among the *TLO* mutants compared to WT, overlap between *TLO* mutants and tested phenotypes, and positive correlation between the number of mutant phenotypes and the number of nucleotide and amino acid changes.

Construction of a PCA biplot for each phenotype and *TLO* mutant highlighted similarities in strain behavior that did not segregate by *TLO* architectural group (Jolliffe and Cadima 2016). The *tlo*Δ/Δ mutant strongly influenced PC1 and displayed the greatest separation from all other *TLO* mutants, likely because of its pleotropic phenotypes that were distinct from single *TLO* mutants. Representatives from both the *TLO*α and *TLO*γ architectural groups were intermixed along PC2 (Fig 5B). Many of the phenotypes that contribute to PC1 are growth related, while stress conditions defined PC2. These stress-related phenotypes provide better separation of individual *TLO* mutants on PC2 than the growth phenotypes associated with PC1.

We hypothesized that *TLO*s with greater sequence divergence from other paralogs may have a proportional decrease in functional redundancy. To quantify this relationship, we constructed phylogenies for the *TLO*α and *TLO*γ architectural group genes to determine their relatedness. For each node in the phylogeny, we calculated the number of nucleotide or amino acid changes and quantifiable phenotypes that distinguished strains or groups of strains on each branch in the phylogeny (Shank, et al. 2018; Madeira, Madhusoodanan, Lee, Eusebi, Niewielska, Tivey, Lopez, et al. 2024; Madeira, Madhusoodanan, Lee, Eusebi, Niewielska, Tivey, Meacham, et al. 2024). A significant positive correlation existed between the number of nucleotide variants or amino acid substitutions and the number of phenotypes observed in deletion mutants for genes at each branch point in the phylogeny (Fig 5C, D, S6). This suggests that an increase in sequence divergence of *TLO* paralogs leads to lower redundancy with other gene family members even though architectural group identity for disrupted *TLO*s was rarely associated with specific phenotypes.

### Loss of single *TLO* genes dysregulates a widely variable number of genes

The role of Tlos as Mediator subunits implies that mutant phenotypes produced by loss of single paralogs occurs through altered gene expression. We measured transcript abundance of the WT, *tlo*Δ/Δ, and each single *TLO* mutant strain during logarithmic phase growth in liquid YPD medium at 30°C to assess molecular redundancy in transcriptional regulation. All single *TLO* mutants displayed changes in transcript abundance despite a lack of any quantifiable difference in growth under these conditions (Fig 1C, 6A, S7). Transcriptional profiles of single mutants formed clusters that included members of multiple architectural groups. For example, the *tlo*α*9*Δ, *tlo*α*12*Δ/Δ, and *tlo*γ*13*Δ/Δ mutants expressed blocks of genes distinctly from all other strains. The remaining single *TLO* mutants have similar trends in gene expression and are distinct from the *tlo*Δ/Δ mutant (Fig 6A).

**Fig. 6:**
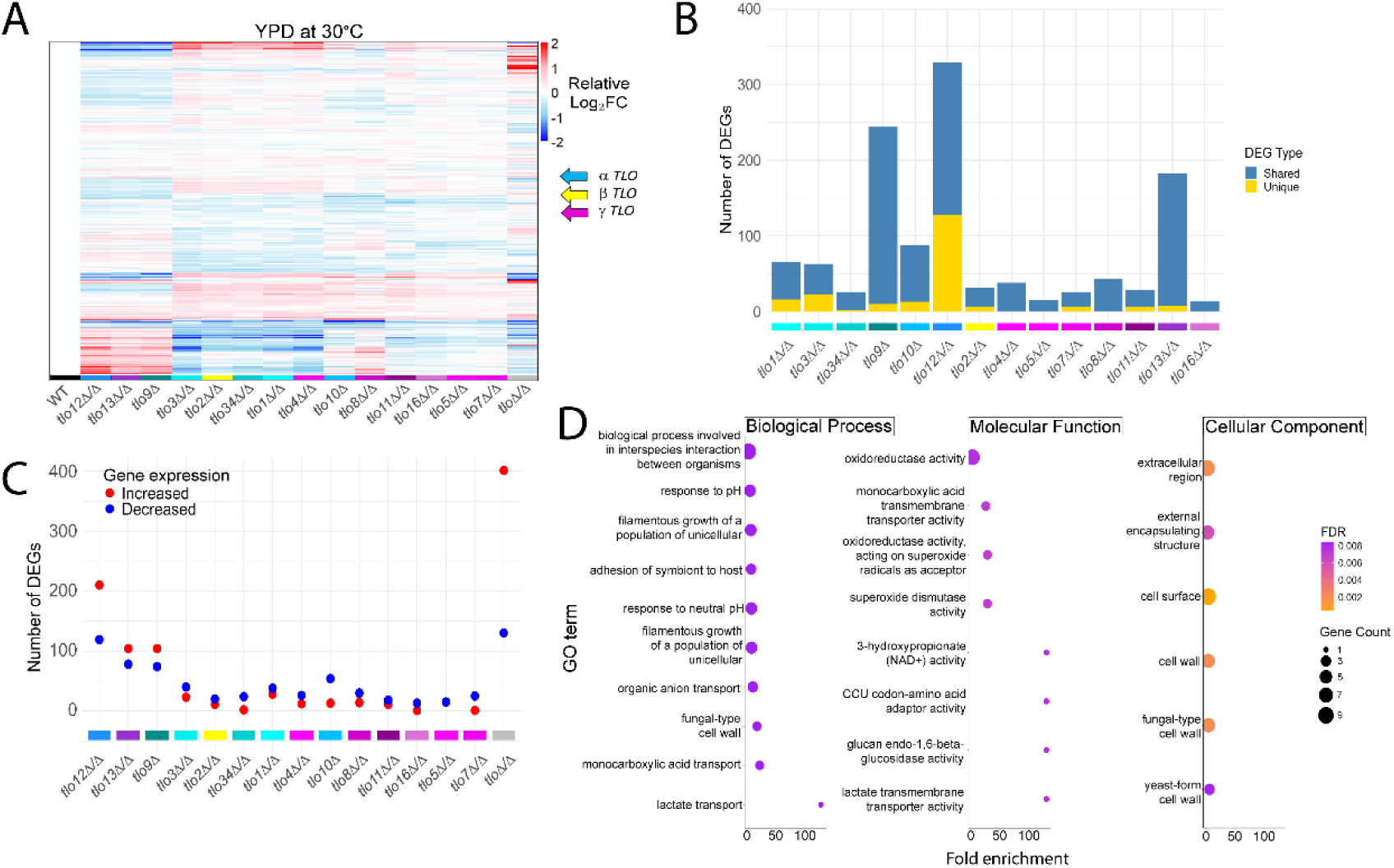
Most genes altered by loss of single *TLO* genes are shared between mutants. (**A**) Differentially expressed genes (DEGs) were identified relative to the wildtype (WT) strain (q≤0.05, ≥2-fold change) for all indicated *TLO* mutants. Expression is color-coded by the relative log_2_ fold change, and *TLO* genes are color-coded by architectural group. (**B**) The number of DEGs were plotted for single *TLO* mutant relative to WT. “Shared” in blue indicates the DEG is also differentially expressed in another *TLO* mutant, and “Unique” in yellow indicates the gene was differentially expressed in a single *TLO* mutant. (**C**) *TLO* mutant strains were ordered by the clustered global transcriptional profiles shown by the heatmap in panel A. The number of DEGs with increased and decreased expression are indicated in red and blue, respectively. (**D**) Shared DEGs present in comparison of ≥7 single *TLO* mutants to WT were annotated for enrichment by gene ontology (GO). A subset of GO categories with the greatest statistical significance are shown and color-coded by their FDR value, with circle size indicating number of genes mapped to that GO term. Significant enrichment was determined by a p-value < 0.01 and corrected by Benjamini-Hochberg method. **Fig. 6 alt text:** Graphs and data of global gene expression profiles and enriched functions that differ between *TLO* mutants and the WT strain, with subfigures labelled a to d.

Single *TLO* mutants had a wide range of differentially expressed genes (DEGs) when compared to the WT (Fig 6B, C). Loss of *TLO*α*12* altered expression of over 300 genes (≥2-fold change, q≤0.05), whereas only 14 genes changed expression in the *tlo*γ*16*Δ/Δ mutant. Yet, most *TLO* mutant strains (12/14), even those showing relatively few genes with altered expression, included unique DEGs not found in any other mutant (Fig 6B). For the three single *TLO* mutants with the greatest number of differentially expressed genes, transcript abundance was more frequently increased compared to those strains with fewer transcriptional changes that had a greater number of genes with reduced expression (Fig 6C). These trends coincide with the general patterns of gene expression in the panel of single *TLO* mutants (Fig 6A).

We explored the genes with altered transcript abundance in at least half of the *TLO* mutants to gain insight into biological, molecular, and cellular processes broadly affected by loss of single *TLO* paralogs. A disproportionate number of differentially expressed genes had roles in filamentation, adhesion, redox reactions, and transporter activity at the cell surface (Fig 6D). Altered expression of genes related to the cell wall structure and transportation of various ion and carbon sources may reflect the prominence of altered filamentation and growth phenotypes among single *TLO* mutants (Fig 5A).

### *TLO* paralogs display synergistic interactions

As transcriptional regulators, Tlo proteins may control expression of other gene family members or coordinately regulate expression of target genes and phenotypes. To assess synergy between *TLO* genes, we constructed two double mutants (*tlo*α*34*Δ/Δ *tlo*β*2*Δ/Δ and *tlo*α*9*Δ/*tlo*α*10*Δ) by iterative rounds of CRISPR/Cas9-mediated *TLO* disruption and compared them to their single *TLO* gene deletion counterparts. Double mutants were selected for construction based on the phenotypes observed in single *TLO* mutants to test potential genetic interactions. The mutant triads (two single mutants and the double mutant) showed additive and synergistic interactions that were dependent on the experimental context (Fig 7A-F). For example, altered growth rates of the *tlo*α*34*Δ/Δ *tlo*β*2*Δ/Δ double mutant matched the additive contributions of each of the single *TLO* mutants under multiple conditions (YPD at 37°C or low phosphate, Fig 7G). In contrast, the *tlo*α*9*Δ/*tlo*α*10*Δ double mutant displayed significantly slower growth in YPM at 30°C than the combined growth delays of the *tlo*α*9*Δ or *tlo*α*10*Δ single mutants. Similarly, loss of either *TLO*α*34* or *TLO*β*2* increased virulence in *G. mellonella* models of infection, but the increased virulence was completely reversed in the *tlo*α*34*Δ/Δ *tlo*β*2*Δ/Δ double mutant. In contrast, the virulence of the *tlo*α*9*Δ/*tlo*α*10*Δ double mutant appeared additive to each of the single *TLO* mutants. Both positive and negative synergy occurred during macrophage killing for the *tlo*α*34*Δ/Δ *tlo*β*2*Δ/Δ and the *tlo*α*9*Δ/*tlo*α*10*Δ double mutant, respectively. In total, double mutants displayed synergy in 10 of 22 assays (Fig 7G). Negative interactions occurred for double mutants in six assays, and positive synergy arose for double mutants in four assays. This indicates that *TLO* genes likely control phenotypes through intersecting or coordinated regulation.

**Fig. 7:**
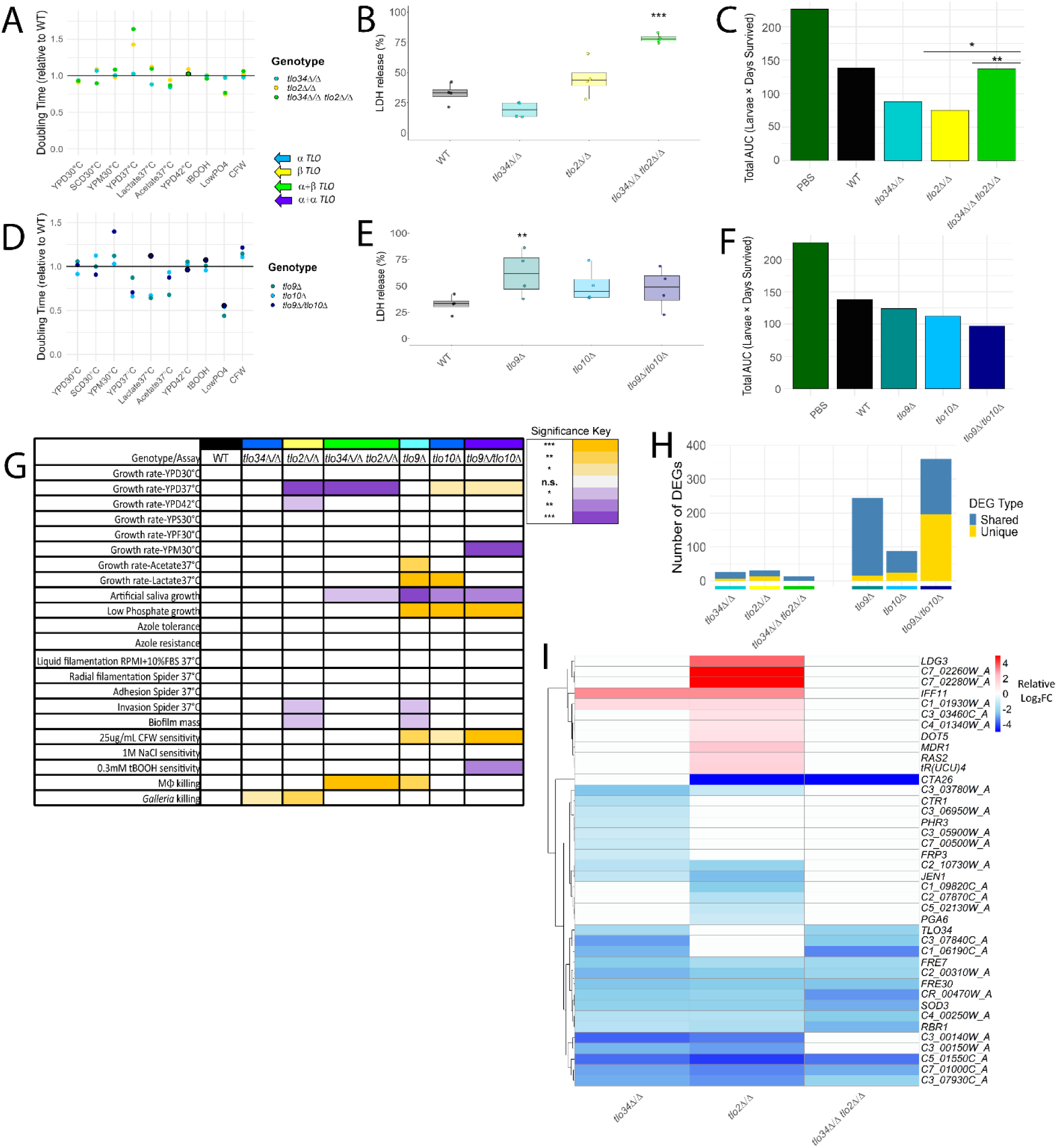
*TLO* mutants display synergistic interactions. (**A, D**) Growth of two double mutants (*tlo*α*34*Δ/Δ *tlo*β*2*Δ/Δ and *tlo*α*9*Δ/*tlo*α*10*Δ) and their corresponding single *TLO* mutants in in media with a variety of carbon sources (YPD/SCD=dextrose, YPM=maltose, lactate, or acetate), temperatures, oxidative stress (tBOOH), low phosphate (0.05 mM PO_4_), and cell wall damage by calcofluor white (CFW). The doubling times were calculated and normalized to the mean WT growth. Each point represents the mean for the color-coded genotype. Significance is indicated by bold outlining and calculated as deviation from an additive phenotype by paired t-tests at p < 0.05. N=4. (**B, E**) RAW 264.7 murine macrophages were incubated with *C. albicans* cells at an MOI of 2. After 20 hours, supernatants were collected, and lactate dehydrogenase (LDH) release was measured. LDH release is plotted for the indicated strains normalized to negative and positive cell lysis controls as the minimum and maximum LDH release. N=4. (**C, F**) *Galleria mellonella* survival is plotted as the area under the Kaplan Meier curves of all animals following infection with 2.5×10^5^ cells for the indicated *C. albicans* strain or PBS as a negative control. Significance was determined via a Log-Rank test with p-value corrections performed by Benjamini-Hochberg method. N=40 animals. (**G**) Double *TLO* mutants and their single mutant counterparts are shown with their relative phenotypic differences compared to WT. White indicates no significant difference from WT, whereas increasing or decreasing phenotypes are indicated by increasing shades of orange or purple, respectively. (**H**) The number of DEGs were plotted for each double *TLO* mutant and their respective single *TLO* mutants relative to WT. “Shared” in blue indicates the DEG is also differentially expressed in another *TLO* mutant, and “Unique” in yellow indicates the gene was differentially expressed in a single *TLO* mutant. (**I**) Heatmap of differentially expressed genes (q≤0.05, ≥2-fold change) for the *tlo*β*2*Δ/Δ, *tlo*α*34*Δ/Δ, and *tlo*α*34*Δ/Δ *tlo*β*2*Δ/Δ mutants relative to wildtype and color-coded by relative log_2_ fold change. Significance was determined via modified Student’s t-tests unless otherwise stated; * p < 0.05, ** p < 0.01, *** p < 0.001. **Fig. 7 alt text:** Graphics and data show the relationships between single *TLO* mutants and their corresponding double knockouts in a series of panels describing phenotypic assays, with subfigures labeled a to g. Molecular phenotypes from differential gene expression analysis of these same strains are shown in subfigures labelled h and i.

Analysis of gene expression in the double mutants further supported synergy between *TLO* genes. The *tlo*α*9*Δ/*tlo*α*10*Δ double mutant had 196 DEGs not present in either single mutant, indicative of positive synergy. In contrast, the *tlo*α*34*Δ/Δ *tlo*β*2*Δ/Δ double mutant differentially expressed a total of 14 genes ((≥2-fold change, q≤0.05), which was fewer than the genes differentially expressed by either single mutant (Fig 7H, 26 and 31 for *TLO*α*34* or *TLO*β*2*, respectively). Individual genes showed instances of conserved expression changes in single mutants and the *tlo*α*34*Δ/Δ *tlo*β*2*Δ/Δ double mutant (*e.g., FRE7*), instances of altered expression in a single mutant but not the double mutant (*e.g., CTR1*), and instances of expression changes in both single mutants that are absent in the double mutant (*e.g., JEN1*, Fig 7I), which highlights the frequent negative synergy observed in the *tlo*α*34*Δ/Δ *tlo*β*2*Δ/Δ double mutant. Taken together, synergistic interactions between *TLO* genes exists at both the molecular and phenotypic levels.

### Regulatory networks connect gene expression and phenotypes of *TLO* mutants

Gene regulatory networks form through the complex interplay of transcriptional regulators to maintain critical cellular functions and coordinate cellular responses. To discern the regulatory networks perturbed by loss of *TLO* genes, we constructed correlation networks using the expression profiles of all single and double *TLO* mutants and the wildtype background strain. We incorporated 3,963 of 6,461 *C. albicans* genes into 32 co-expression modules that varied in size from 895 to 25 genes (Fig 8A, S8). All co-expression modules, except for the magenta module, were significantly correlated with at least one *TLO* mutant genotype (Fig S8). A single module was correlated with mutants either lacking *TLO*β*2* or *TLO*γ*7*. Interestingly, the same 8 modules were associated with the loss of *TLO*α*3*, *TLO*α*9*, *TLO*α*12*, and *TLO*γ*13,* but the directionality of these correlations was inverted between *tlo*α*3*Δ/Δ and the other mutants (Fig S8). Consistent with a larger impact on expression, the greatest number of modules were associated with the *tlo*α*12*Δ/Δ and the *tlo*α*9*Δ/*tlo*α*10*Δ mutants (19 of 32 modules), three of which showed extremely strong associations (*e.g.,* brown, yellow, and orange modules, Fig S8).

**Fig. 8:**
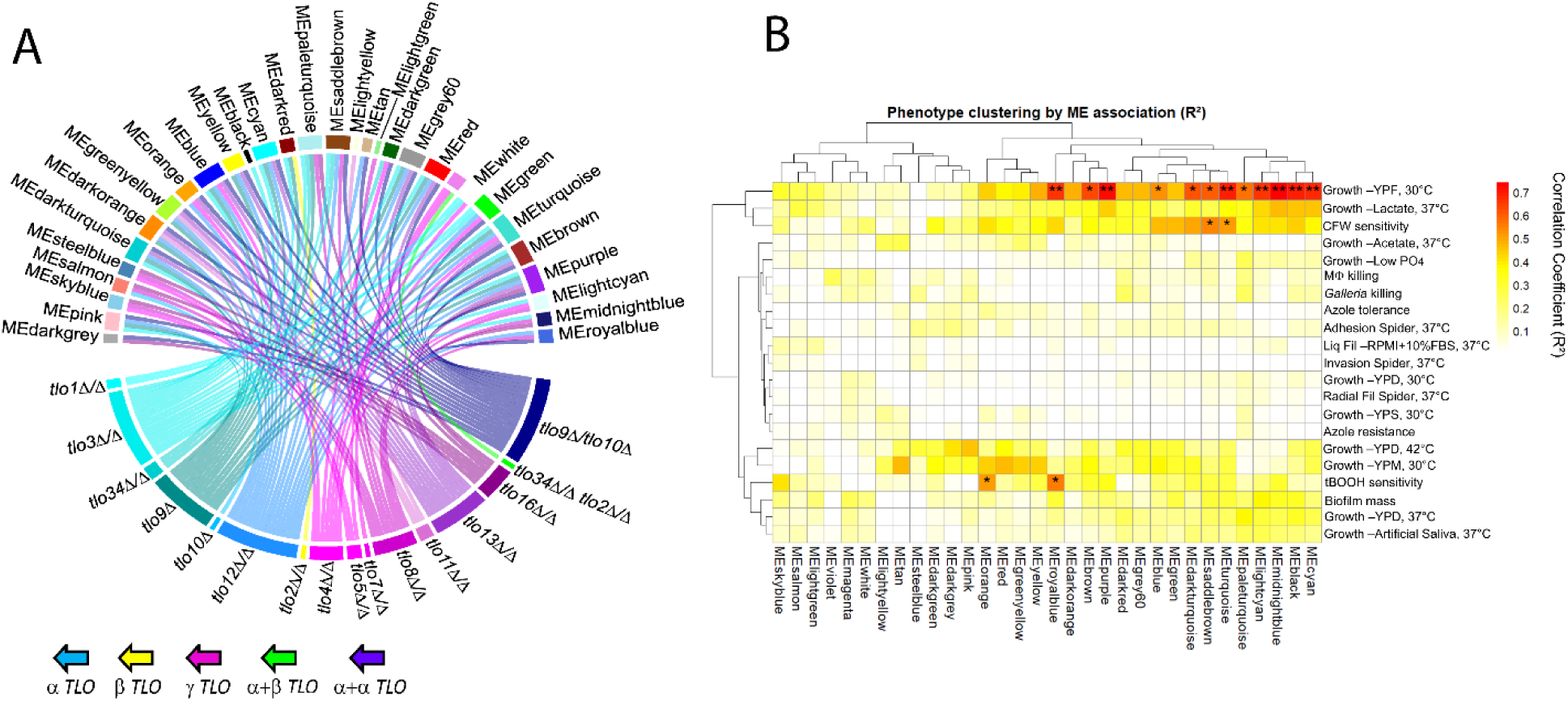
Identification of gene co-expression module and phenotype connections revealed by loss of *TLO* genes. (**A**) A chord diagram connects significant interactions between *TLO* mutants and the 32 constructed co-expression modules. (**B)** The thirty-two module eigengenes (MEs) were correlated against the raw phenotype data for all strains in each assay by Pearson correlation. * denotes p < 0.05. ** denotes p<0.01. **Fig. 8 alt text:** Graphics detail the relationships between *TLO* mutants and gene co-expression modules from WGCNA analysis and associated mutant phenotypes, with subfigures labeled a and b.

Module-phenotype correlations pointed to genes potentially responsible for phenotypic response among the *TLO* mutants. For example, the turquoise module connected decreased growth rates in fructose and CFW with gene expression in the *tlo*α*9*Δ, *tlo*α*12*Δ/Δ, *tlo*γ*13*Δ/Δ, and *tlo*α*9*Δ/*tlo*α*10*Δ mutants (Fig 8B, S8). The increased sensitivity of these mutants may be linked to the decreased transcript abundance of several genes (*e.g., PDR16*, *HSP70*, and *HSP12*) that are important for resistance to chemical stressors (Smith, et al. 2004; Liu, et al. 2005; Enjalbert, et al. 2006; Saidane, et al. 2006).

### *TLO* gene family expression does not associate with mutant phenotypes

To investigate if altered expression of the *TLO* gene family contributes to the mutant phenotypes, we compared the expression of each *TLO* gene and total *TLO* transcript among the constructed mutants. *TLO*α genes had higher transcript levels than their *TLO*γ counterparts, and the *TLO*β2 member was intermediate in expression between the two groups (Fig S9A), consistent with previous reports (Anderson, et al. 2012; Zhang, et al. 2012). *TLO* genes displayed a similar pattern of expression across all mutants although the precise level of expression fluctuated slightly among the mutant panel. As expected, expression of the deleted *TLO* gene was dramatically reduced in each mutant although many mutants retained some low level of expression that is likely a consequence of multi-mapped reads being randomly assigned to one of the multiple remaining paralogs. Comparison of total *TLO* gene expression among the single *TLO* mutants did not find significant differences when compared to the wildtype strains or other *TLO* mutants, with the exception of slightly elevated *TLO* expression in *tlo*γ*16*Δ/Δ (Fig S9B). In contrast, loss of two or more *TLO* genes led to a reduction in gene family expression. Total *TLO* expression was reduced in the *tlo*α*9*Δ/*tlo*α*10*Δ double mutant compared to the wildtype and was undetectable in the complete *tlo*Δ/Δ mutant. This suggests that *TLO* expression can be influenced by the loss of paralogs but is unlikely to be responsible for the phenotypic variation observed in the single *TLO* mutant panel.

## DISCUSSION

Repeated gene duplication is a critical process to the production of gene repertoires that can improve organismal fitness. This investigation of the *C. albicans TLO* expanded gene family supports a model by which closely related paralogs retain a strong functional overlap with other gene family members while also acquiring non-redundant functions that are detected in ∼20% of the total strain x phenotype assays. Assignment of *TLO* paralogs to an architectural group did not influence the degree of redundant and non-redundant function, which ranged widely among single *TLO* deletion mutants. Instead, the number of new sequence variants at each branchpoint in the *TLO* group phylogeny more strongly predicted the extent of non-redundant phenotypes in mutants for single *TLO* genes. Furthermore, altered phenotypes were more commonly observed for single *TLO* deletion strains under stressful conditions and were often observed for multiple single *TLO* mutants. Altered expression of the same genes was also commonly found among single *TLO* mutants, suggesting that redundancy may arise from coordinated regulation of target genes by multiple Tlos or the presence of feedback loops among *TLO* genes. These interconnected regulatory relationships could protect cells from the detrimental consequences of single gene loss in rapidly evolving gene families.

This study systematically addresses longstanding questions of redundancy and functional divergence among individual paralogs following the formation of gene families. Overall, most *TLO*s displayed substantial molecular and biological overlap with other gene family members, which might be expected for recent lineage-specific expansions. Yet, every *TLO* paralog was associated with mutant phenotypes, indicative of non-redundant functions. Interestingly, stressful conditions (*e.g.,* heat shock, cell wall stress, low phosphate) produced observable phenotypes more frequently than nutrient replete conditions and may indicate that *TLO* expansion promoted stress survival. We posit that the presence of observable phenotypes in single *TLO* mutants suggests that the some degree of subfunctionalization or specialization for minor functions occurred during expansion (Fig 9). Multiple *TLO* mutants displaying altered phenotypes under the same set of conditions argues for a more conserved original function that was partitioned among paralogs during expansion as opposed to repeated neofunctionalization. Subfunctionalization could arise through the binding of Tlos to different genes within the same pathway, leading to similar phenotypic outcomes (Nobile, et al. 2012; Hernday, et al. 2013). Alternatively, crosstalk among *TLO* gene family members could reproduce our results. Tlo proteins may regulate other *TLO* genes directly as they have been shown to play a role in subtelomeric gene expression in *S. cerevisiae* (Peng and Zhou 2012), indirectly through other transcription factors altered in multiple *TLO* mutants (*e.g.*, *FCR1*, *MSN4*, *UME6*, etc.), or bind to the same promoters to coordinate gene expression. Evidence for coordinated regulation of *TLO* genes is supported by the high sequence identity in the upstream and downstream regulatory sequences flanking the coding sequence. Loss of a paralog could increase regulation of the remaining paralogs by upstream transcriptional regulators. Tlo proteins acting at the same promoter in target genes could also reproduce the observed antagonistic effects in *TLO* mutants. One Tlo may repress expression, whereas another Tlo activates expression. Loss of either *TLO* gene would upset this balance and lead to altered expression.

**Fig. 9:**
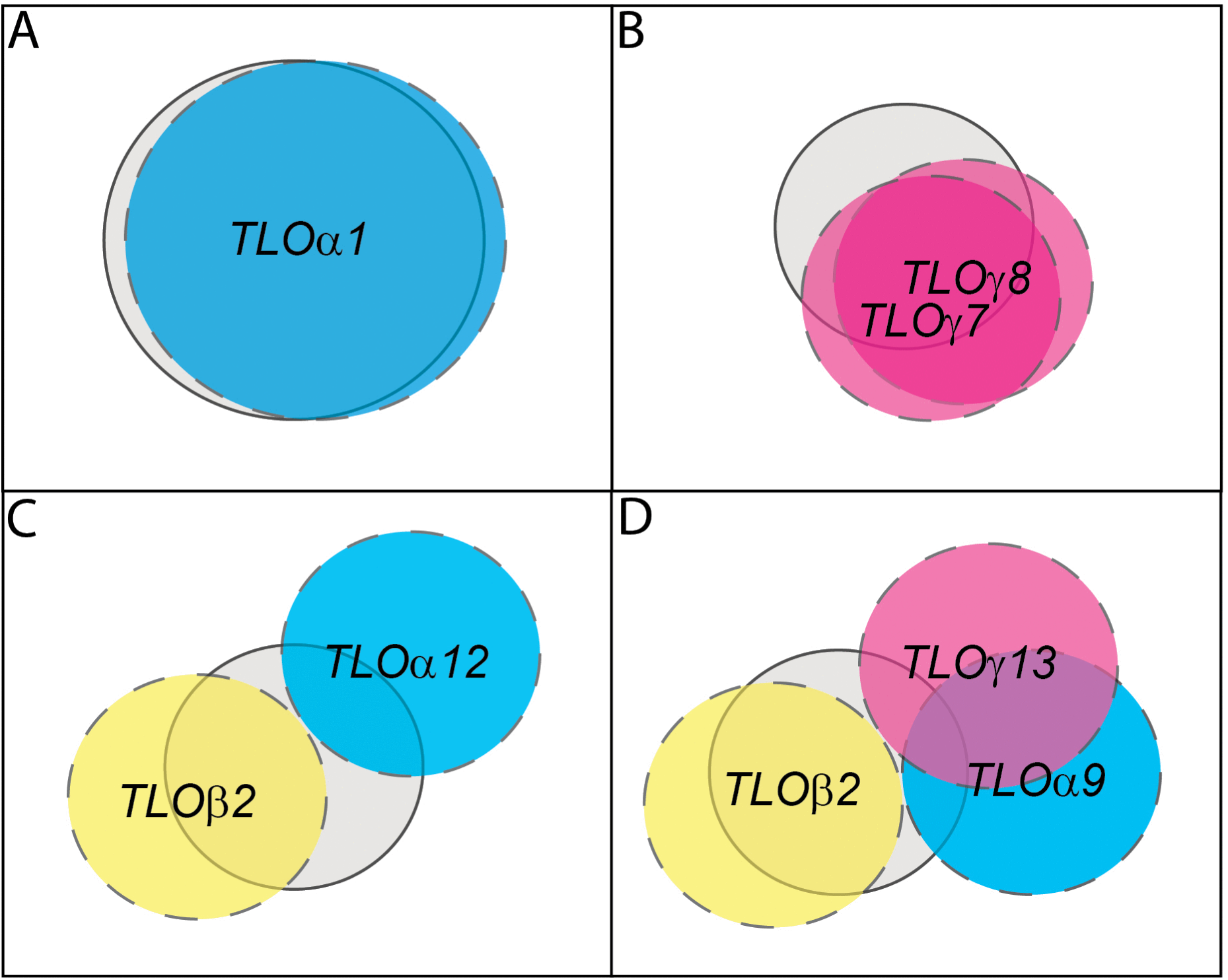
Model of functional redundancy among *TLO* paralogs. *TLO* genes exhibit various degrees of functional redundancy with other family members. (**A**) Loss of *TLO*α*1* produced very few phenotypic changes, suggestive of significant functional overlap with other *TLO* family members (grey circle). (**B**) *TLO*γ*7* and *TLO*γ*8* share multiple phenotypes when disrupted but display some amount of unique phenotypes compared to each other. (**C**) *TLO* genes can display completely independent sets of phenotypes when disrupted as is observed for *TLO*α*12* (cyan) and *TLO*β*2* (yellow). (**D**) The partial similarity of the phenotypic profile for the *tlo*α*9*Δ mutant is overlaid on the *tlo*α*12*Δ/Δ mutant but has no functional overlap with *tlo*β*2*Δ/Δ. **Fig. 9 alt text:** Graphical summary of representative TLO paralog functions overlapping with putative functional space of the ancestral gene, with subfigures labelled a to d.

Sequence divergence between *TLO*s was the strongest predictor of non-redundant phenotypes among *TLO* mutants. Extensive characterization of the outcomes of gene duplication have been used to describe how functional divergence arises through mutation to often produce paralogs with partially overlapping functions (Huang and Golding 2012; Haran, et al. 2014; Laurent, et al. 2020; Tang, et al. 2024; Dennler and Ryan 2025). Investigation of all paralogs from the *TLO* gene family here demonstrated that the presence of multiple other paralogs did not constrain non-redundant functions. Additionally, no obvious relationship between the frequency of non-redundant functions and the predicted chronology of *TLO* emergence (*TLO*β*2* ◊ *TLO*α ◊ *TLO*γ (Jackson, et al. 2009; Anderson, et al. 2012; Dunn, et al. 2022)) was evident. The absence of an architectural influence on gene function is supported by prior work on *TLO* genes (Dunn, Kinney, et al. 2018), but we cannot rule out that genetic drift could lead to similar phenotypic profiles in single *TLO* mutants from different architectural groups. Expression of *TLO* genes was not substantially altered by loss of a single paralog when grown in rich medium, suggesting that expression changes among paralogs in single *TLO* mutants is unlikely to be responsible for the observed phenotypes. Additionally, stronger or more frequent phenotypes were not observed in mutants for the more highly expressed *TLO*α genes. Therefore, function and redundancy among *TLO* members currently appears to be strongly influenced by the accumulation of small changes in sequence.

A major strength of investigating paralog redundancy with the *TLO* genes is their pleiotropic function as transcriptional regulators. Most investigations of paralog redundancy have focused on genes encoding effectors that function in a single biological process. Measuring a single phenotype limits the ability to observe effects caused by loss of a single paralog (Carlisle, et al. 2013; Ewen-Campen, et al. 2017; Kwon, et al. 2022). In contrast, the expansion of *TLO* genes and other transcriptional regulators can impact the expression of many genes with broad roles in organismal physiology (Myers, et al. 1999; Ansari, et al. 2012; Haran, et al. 2014; Dolan, et al. 2017). This likely aided our ability to observe minor changes in gene expression and infrequent phenotypic changes in mutants for single *TLO* genes (*e.g.*, *TLO*α*34*) to identify instances of non-redundant function. Surprisingly, the direction of phenotypic changes associated with loss of single *TLO* genes was often inverted in the *tlo*Δ/Δ mutant. This suggests that misleading definitions of the function of each paralog can be constructed based on inhibition of all gene family members as more complex regulatory or functional relationships might be overlooked.

Phenotypes caused by the loss of single *TLO* genes may indicate that Tlo proteins are sensitive to perturbation. The available pool of Tlo protein exceeds the stoichiometric requirements as subunits of the Mediator complex (Zhang, et al. 2012; Liu, et al. 2016), and would be expected to be unaffected by loss of a single paralog, especially the more lowly expressed *TLO*γ genes (Anderson, et al. 2012). So, how does loss of a single isoform of an interchangeable protein subunit cause such dramatic changes in cell behavior? The loss of a single *TLO* gene could alter accessibility of other paralogs to Mediator and change the functional output of the cell. Thus, loss of a specific gene (paralog A) could directly reduce a transcriptional response and loss of a separate gene (paralog B) could increase the access of paralog A to Mediator and increase a transcriptional response. This would provide another possible explanation for bidirectional phenotypic changes among *TLO* mutants. Alternatively, phenotypic alterations caused by loss of single *TLO* genes may suggest that they can act independently of Mediator through a previously described C-terminal transcriptional activation domain (Liu and Myers 2015). This is further supported by the equal frequency of *TLO*γ mutants producing altered phenotypes as the other architectural groups even though biochemical analysis of Mediator failed to identify any associated Tloγ proteins (Zhang, et al. 2012). Indeed, it is tempting to speculate that the prior localization of Tloγ-GFP fusion proteins to the mitochondria (Anderson, et al. 2012) may indicate that mitochondrial gene expression is regulated by the Tloγ architectural group. Alternatively, Tlos could regulate nuclear gene expression independent of Mediator as part of subcomplexes, as has been described in *Saccharomyces cerevisiae* (Peng and Zhou 2012), or manipulate protein-protein interactions (Zhu, et al. 2015; Liu, et al. 2016), including those of other Tlos and their association with Mediator. Function outside of Mediator could therefore explain phenotypes arising in mutants lacking single *TLO* genes and explain, in part, why they are present in stoichiometric excess of Mediator.

Analysis of *TLO* double mutants highlights the ability for paralogs of the same gene family to form complex relationships. With only two double mutants, we observed evidence of positive synergy, antagonism, and a lack of interaction between paralogs for different phenotypes. No bias towards an interaction type was evident from the data collected here but is constrained by the use of only two double mutants. Regardless, our data supports the existence of complex interactions among paralogs that was only inferred from analysis of the single *TLO* mutants. Of note, loss of two genes from different architectural groups produced fewer phenotypes than loss of two *TLO*α genes, which may indicate stronger redundancy existing among closely related sequences.

The landscape of redundancy for *TLO* genes serves as a useful model for other gene families that have undergone lineage-specific expansions. Like many other gene families, the *TLO*s exist in a repetitive region of the genome that allows for rapid copy number changes over short evolutionary timescales. Therefore, we speculate that as gene families expand, we might expect that the functional space for a new paralog to occupy would shrink, limiting the likelihood for new genes to be retained when they arise. However, substantial functional overlap among the *TLO*s suggests that a small amount of unique or coordinated function for a gene family member may be sufficient to overcome the potential fitness costs of maintaining additional paralogs. This may give enough functional space for drift or selection to confer new roles to duplicated genes. Additionally, functional redundancy or overlap among large paralogous gene families might provide a selective advantage by safeguarding advantageous functions in those genes that undergo frequent mutation in highly plastic regions of the genome. If those gene functions are critical for organismal survival, this would allow mutation to produce new alleles to be tested for fitness without posing a significant cost. Finally, the environmental context of the organism needs to be considered when interpreting the outcomes of paralogs. Stable environments could select for beneficial alleles prior to duplication and reduce any fitness gains from gene duplications. In contrast, fluctuating and disparate environments, such as those experienced by *C. albicans* during colonization of the complete human host, could favor repeated rounds of innovation and retention of new paralogs that may shift the composition of the gene family over time. This could favor combinations of redundant and unique function among gene family paralogs as a bet-hedging mechanism to support long-term survival in the host.

## METHODS

### Media and reagents

Yeast extract-peptone-dextrose (YPD), synthetic complete dextrose (SCD), and Spider media were prepared as previously described (Chauhan and Kruppa 2009; Dymond 2013). YPD plates for selection were prepared by adding nourseothricin (NAT) at a final concentration of 200 μg/mL (Jena Bioscience, Jena, Germany). Growth in maltose, fructose, or sucrose was assessed by replacing dextrose with the specified carbon source in the YPD recipe at a final concentration of 2%. YPD medium was supplemented with 1 M NaCl or 25 μg/mL of calcofluor white (CFW) for high salt and cell wall stress media, respectively. Modified SCD recipes were used for assaying acetate and lactate by replacing dextrose with 2% sodium acetate or sodium L-lactate to a final concentration of 0.4%. Low phosphate medium was prepared as previously described and supplemented with 1 M KH_2_PO_4_ to a final concentration of 0.5 mM (Acosta-Zaldivar, et al. 2024). Artificial saliva medium was made with sterile synthetic saliva (Millipore Sigma, Burlington, MA, USA) and supplemented with 0.005 mg/mL glucose and 1.19 mg/mL porcine gastric mucin (Millipore Sigma, Burlington, MA, USA) to mimic the major abiotic factors within the human oral cavity (Valentijn-Benz, et al. 2015).

All strains were struck from –80°C stocks onto YPD agar medium and used for assays within 2 weeks. Two independent lineages were used for all experiments. Unless otherwise noted, strains were incubated in 2 mL YPD overnight at 30°C prior to use and experiments were conducted with 4 biological replicates (two replicates for each of two genetic lineages).

### Strain construction

All strains, oligonucleotides, and plasmids used in this work are provided in Supplementary Tables S1-S3. *TLO* mutant strains were constructed from a *LEU2* heterozygous SC5314 strain using a modified *C. albicans* CRISPR/Cas9 protocol (Nguyen, et al. 2017). Briefly, guide RNAs (gRNAs) were designed in the Benchling software to maximize on-target scores and minimize off-target scores to the *TLO* locus of interest (https://www.benchling.com). All gRNAs corresponded to sequences within 2 kb of the target open reading frame (ORF) to initiate loss of the *TLO* ORF. Clean gene deletions were produced by designing repair template donor DNA (dDNA) that aligned to intergenic regions flanking the gRNA target site and the *TLO* ORF.

Transformations were performed using a modified lithium acetate with heat shock approach. The transformant DNA components were prepared as in Nguyen *et al*. and added to a master mix of 1 M lithium acetate, 10X TE, 5 mg/mL boiled/sheared salmon sperm DNA, 1 M DTT in H_2_O, and 50% PEG 3350 (Nguyen, et al. 2017). This was added to approximately 2.0×10^8^ *C. albicans* cells taken from a saturated culture grown overnight in YPD at 30°C. Cell mixtures were incubated at 30°C for 1 hr and were exposed to a heat shock at 42°C for 45 minutes (min). Cells were then centrifuged at 2200 revolutions per minute (rpm) for 5 min, resuspended in sterile H_2_O, and plated directly to YPD+NAT selection plates. Plates were incubated at 30°C for 1-3 days until putative transformant colonies were big enough to patch out and validate by PCR. Successful colonies were plated to SCD lacking L-leucine to remove the CRISPR cassette that contains Cas9 during repair of the single copy of *LEU2*, as described by Nguyen *et al*. (Nguyen, et al. 2017).

Following selection, colonies were checked twice for loss of the target locus via cell crush PCR using the commercial polymerase Accustart II (Quantabio, Beverly, MA, USA). Colonies containing the desired mutation had genomic DNA (gDNA) extracted using the Zymo Quick-DNA Miniprep kit (Zymo Research, Irvine, CA, USA). DNA concentration was established using the Qubit 3 Fluorometer with the dsDNA Broad Range Assay Kit (Thermo Fisher Scientific, Waltham, MA, USA). Finally, gDNA libraries were built using the NEBNext Ultra II FS DNA library prep kit for Illumina (New England Biolabs, Ipswich, MA, USA) prior to whole genome sequencing (WGS) or submitted as gDNA for preparation by The Ohio State University Applied Microbiology Services Laboratory (AMSL)) or the University of Wisconsin Biotechnology Center DNA Sequencing Facility (UWBC). AMSL utilized the Illumina NextSeq 2000, while UWBC processed samples on the Illumina NovaSeq X Plus. Raw sequencing data was processed in command line and quality assessed via FastQC analysis (Andrews 2010). Trimmomatic was used to remove any low quality sequences and remaining adapter sequences with the following thresholds: SLIDINGWINDOW:4:20 MAXINFO:125:1 LEADING:20 TRAILING:20 HEADCROP:20 MINLEN:35 (Bolger, et al. 2014). Bowtie2 was used to align samples to the *Candida albicans* SC5314 Ca21 reference genome assembly or an in-house phased SC5314 long read genome assembly (Langmead and Salzberg 2012). Samtools software was used to convert.sam files into.bam format, sort the files, and index them (Danecek, et al. 2021). Finally, processed files were inspected manually for successful deletion of the target locus in the Integrated Genome Viewer (IGV) software (Robinson, et al. 2011). To ensure that strains were free of aneuploidies or long tract loss of heterozygosity (LOH) events, WGS data was visualized compared to the Ca21 reference genome with YMAP (Abbey, et al. 2014).

### Growth rates

Growth assays were conducted as described previously (Woodruff, et al. 2024). Briefly, overnight cultures were grown in 300 µl of liquid YPD medium at 30 °C in a 96 deep-well plate, with shaking at 125 rpm. The following day, overnight cultures were diluted 1:40 into H_2_O and then 1:50 into fresh liquid growth medium into a final volume of 150 µl in a clear Greiner CELLSTAR 96-well flat-bottom cell culture plate (Greiner Bio-One). The plate was then sealed with a sterile, optically transparent polyester adhesive sealing film. Optical density at 600 nm (OD_600_) was measured every 15 min. for 18-72 hrs, or until growth saturation was reached, at specified temperatures using a BioTek Synergy H1 microplate reader (BioTek Instruments, Winooski, VT, USA) on double orbital continuous shaking at fast orbital speed and a frequency of 425 cycles per minute (cpm). The minimum doubling time was calculated using the polynomial measurement of the curve in the *growthcurveR* package (Sprouffske and Wagner 2016). Each growth curve experiment had two technical replicates for each strain.

### Spot dilution assay

Aliquots from saturated overnight cultures were diluted 1:10 in sterile 1X PBS. Cells were counted and diluted to 1×10^7^ cells/mL. Cells were then serially diluted with 1xPBS starting at the 1×10^7^ cells/mL concentration, 1:10, 1:5, 1:5, and 1:5. YPD, YPD+NaCl, and YPD+CFW plates were spotted with 3 μL of cells from each dilution. Plates were dried with the lids off for 15-20 minutes in a pre-sterilized biosafety cabinet and then incubated at 30°C for 48 hrs before images were taken using a Bio-Rad ChemiDoc and Image Lab software (Bio-Rad, Berkeley, CA, USA).

### Azole disk diffusion assay

A NanoDrop One^C^ spectrophotometer was used to measure the OD_600_ of each overnight culture (Thermo Fisher Scientific, Waltham, MA, USA). Cultures were then diluted to an OD of 0.04 and 70 μl were plated on YPD agar using sterile glass beads. Plates were dried for an hour before a sterile disk infused with 25 μg of propiconazole was placed at the center of the plate with sterile tweezers (Millipore Sigma, Burlington, MA, USA). Plates were incubated for 48 hrs at 30°C prior to imaging with a Bio-Rad ChemiDoc. The radius of inhibition (RAD) and fraction of growth (FoG) scores were calculated with the *diskImageR* script (Gerstein, et al. 2016).

### Liquid filamentation

Overnight cultures were centrifuged at 2200 rpm for 3 min, washed with 1xPBS, and resuspended in 1mL of 1xPBS. 20 μl of washed cells were added to 980μl of RPMI + 10% FBS. Cells were incubated at 37°C for 4 hrs at 225 rpm in a shaking incubator. Following incubation, cells were fixed with 1:10 volume of 37% formaldehyde in a roller drum for 15 minutes at room temperature. Fixed cells were centrifuged and resuspended in 1 mL of 1xPBS. To break up adhered clumps of cells, a Fisherbrand probe sonicator (Fisher Scientific, Waltham, MA, USA) was used (40% power for 10 sec and 15 sec of rest repeated 3 times) while cells were kept on ice to prevent overheating. Cells were imaged at 40x magnification using a Leica DM750 compound light microscope, with an attached Leica MC170HD digital camera and LAS V4.12 software (Leica Microsystems, Wetzlar, Germany). At least 50 cells were visualized per replicate across 5 or more complete fields of view.

### Solid filamentation

Strains were assessed for radial filamentation, adhesion, and invasion on Spider agar as previously described with minor alterations to the visual analysis recipes (Dunn, Kinney, et al. 2018; Dunn, et al. 2020). Briefly, cells were counted on a hemacytometer, and 100 cells were plated onto solid Spider medium. Plates were grown at 37°C for 7 days before being imaged in a Bio-Rad Chemidoc. Images were processed and analyzed using the visual analysis software MIPAR (MIPAR, Worthington, OH, USA). Degree of radial filamentation was calculated using the following equation: Rad_fila_ = (area_hyphal growth_ – area_center colonies_) / (area_center colonies_), where area is defined as the summed pixel value of every colony on the plate. Following capture of the first image, the plates were rinsed with a steady stream of water for 3-5 seconds to remove any non-adherent cells. Adhesion was calculated as Ad= (area_colonies post-rinse_)/ (area_colonies pre-rinse_). Finally, colony invasion was detected by washing the surface of the plate under a steady stream of water and rubbing the agar surface with a gloved finger. Plates were allowed to dry for 30-60 minutes, and imaged. Invasion was calculated as the (area_hyphal agar invasion_)/ (area_colonies pre-wash_).

### Biofilm formation

Silicone elastomer sheets (Bentec Medical, Woodland, CA, USA) were cut into 1×1 cm squares, weighed, and sterilized with UV light in a sterile BSC for 15 minutes. Silicon squares were pre-treated with adult bovine serum (ABS) at 37°C overnight in 12-well plates (Fisher Scientific). The next day, squares were washed in 1xPBS before being placed in 2 mL of liquid Spider medium. The treated silicone squares were incubated with 0.5 OD_600_ of cells and incubated at 37°C for 90 minutes at 120 rpm. Non-adherent cells were removed by dipping the silicone squares in sterile 1xPBS, then placed in fresh Spider medium for 66 hrs at 37°C with shaking at 120 rpm. After incubation, the Spider medium was aspirated from each well and the silicone squares were allowed to dry for 24 hrs, and the post-experiment mass was recorded. Experiments used 3 biological replicates for each genotype.

### Macrophage killing

RAW 264.7 macrophages were thawed at room temperature from long-term storage in DMSO at –80°C. Macrophages were spun down for 5 min at 100 x g and DMSO supernatant was removed. The cell pellet was resuspended with 14 mL of pre-warmed DMEM containing L-glutamine mixed with 10% FBS and 1% α/α (penicillin-streptomycin-Gibco Amphotericin B) (Thermo Fisher Scientific, Waltham, MA, USA). The macrophages were seeded into a vented TC-treated T25 flask and incubated at 37°C + 5% CO_2_ and allowed to adhere to the bottom of the flask and grow until 70-80% confluent before passaging to fresh growth medium in a T75 flask. Cells were passaged 14 times prior to use in LDH release assays. RAW cells were detached from TC-treated T75 flasks with trypsin, stained with 1:1 volume of trypan blue to determine percent viability, then seeded in a TC-treated flat-bottom 96-well plate at a concentration of 25,000 cells per well. Following 24 hrs of incubation at 37°C in 5% CO_2_, *Candida albicans* cells were enumerated with an automated cell counter and seeded to each well at an MOI of 2. After 20 hrs of co-incubation, wells were treated with reagents from the Promega CytoTox96^TM^ Nonradioactive Cytotoxicity Assay kit according to instructions (Promega Corporation, Fitchburg, WI, USA). Macrophage killing via lactate dehydrogenase (LDH) release by *TLO* mutants was measured in a Tecan plate reader at OD_490_ (Tecan Life Sciences, Männedorf, Zürich, Switzerland). Each experiment featured a complete lysis well and a no lysis well as positive and negative controls, respectively.

### Galleria mellonella virulence

*Galleria mellonella* larvae were infected with *C. albicans* and assessed for virulence as previously described (Dunn, Woodruff, et al. 2018). Larvae were ordered from speedyworm.com, with overnight shipping (Speedy Worm, Alexandria, MN, USA). Once delivered, all larvae were stored at ambient temperature. Briefly, *C. albicans* cells were washed with 1x PBS three times, counted with a hemacytometer, diluted to 2.5×10^7^ cells/mL in 1x PBS, and 2.5×10^5^ cells injected into larvae in 10 μl of 1xPBS. Prior to each injection, a 26G Hamilton needle and syringe (Hamilton Company, Reno, NV, USA) was washed in 100% EtOH followed by 1xPBS. Control larvae were injected with 10 μl of sterile 1xPBS. A dilution of 100 cells was plated to solid YPD, and CFUs counted to confirm proper inoculum. Larvae were stored in sterile petri dishes with a small amount of sawdust at 37°C and monitored for 7 days post-infection. Survivors were counted each day and dead larvae were removed from the dish. Pupated larvae were removed from the petri dish and final analysis if present, as virulence in this life stage is unknown. Each replicate included 10 larvae.

### Construction of PCA biplot

Phenotypic data for each single *TLO* mutant and the *tlo*Δ/Δ was converted to a signed (positive or negative) significance score using the summary table in Fig5 and columns lacking significant changes from WT were removed. A principal component analysis (PCA) was performed using the remaining values with the prcomp command and the phenotype data was converted into vectors and plotted. The contributions of PC1 and PC2 were obtained by retrieving the top two levels contributing to variance in the data with *stats* (v 4.4.0). To overlay strain genotypes, the data was scaled and coordinates were extracted to plot each genotype in *ggplot2* (v 3.5.1): sqrt(scores$PC1^2 + scores$PC2^2), scores$contribution – min(scores$contribution)) / (max(scores$contribution) – min(scores$contribution))). We then extracted the plot coordinates and computed the squared distance from the origin as “quality of representation” using pca$x[, 1:2], rowSums(ind_coords^2) / rowSums(pca$x^2). This code was generated with assistance from ChatGPT-4 (OpenAI 2023).

### Analysis of sequence divergence versus functional innovation

Nucleotide sequences for the *TLO*α and *TLO*γ architectural group members were aligned using the EMBL-EBI MUSCLE (v 5) Job Dispatcher website under ClustalW output format (default parameter) (Edgar 2004). The alignment was used to produce phylogenies with the integrated phylotree.js software (Shank, et al. 2018). Phylograms were produced using default parameters (distance-matrix UPGMA) and visualized for the α and γ *TLO*s separately. Sequences from each of two branches for each node were compared to identify the number of nucleotide and amino acid sequence differences for all sequences represented by the two branches. Phenotypic differences between the two groups of genes separated by each node were also counted. This was performed for all nodes in each of the *TLO*α and *TLO*γ phylogenies separately. Finally, the mutations (nucleotide variants or amino acid substitutions) were plotted against the number of phenotype differences and a line of best fit was assigned via the linear model function in R. Significance was determined using Pearson’s correlation.

### RNA sequencing (RNA-seq) and differential gene analysis

Four independent 2 mL cultures for each genotype, two replicates from each of two independent lineages for all mutants, were grown overnight in YPD at 30°C, subsequently diluted 1:100 (1:50 for the slow growing *tlo*Δ/Δ strain) into fresh YPD and grown at 30°C for 2 hrs. to reach mid-log phase growth. RNA was extracted and purified using the MasterPure Yeast RNA Extraction Kit (LGC Biosearch Technologies, Hoddesdon, Herts, UK) with minor adaptations to the base protocol; specifically, DNase I (Thermo Fisher Scientific) treatment of RNA was extended to 1 hr to ensure removal of all DNA prior to storage at –80°C. RNA concentration and purity was measured via Nanodrop, and poly(A) RNA was extracted from 1 mg of RNA for each sample with the NEBNext Poly(A) mRNA Isolation Module (NEB #E7490). The purified mRNA was used to construct cDNA libraries with the NEBNext Ultra II Directional RNA Library Prep Kit (NEB #E7760) dUTP synthesis method (Parkhomchuk, et al. 2009). Library quality and fragment distribution was determined using a DNA High Sensitivity Kit on an Agilent 2100 Bioanalyzer (Agilent Technologies, Santa Clara, CA, USA). All libraries were sequenced to a depth of 20 million reads as 150 basepair paired-end reads on an Illumina NovaSeq X Plus.

The resulting FASTQ files were quality controlled via FASTQC. RNA-seq reads were aligned to the indexed SC5314 genome reference assembly, Ca21, using Spliced Transcripts Alignment to a Reference (STAR, v 2.7.11b) (Dobin, et al. 2013). Raw count tables were generated using HTSeq (v1.99.2) (Anders, et al. 2015). From these raw counts, we calculated normalized expression levels as transcripts per million (TPM) for all genes. All RNA-Seq datasets clustered by their genotype.

Differentially expressed genes (DEGs) were identified using *DESeq2* (v 1.99.2), with the WT samples serving as the expression control for all mutant strains (Love, et al. 2014). The initial DEGs (any genes with a log_2_fold change (LFC) > or < 0) were subjected to adaptive shrinkage via the *ashr* package (v 2.2-63) in R (Stephens 2017). Significant DEGs were determined by using cutoff criteria of an adjusted p-value < 0.05 and a 2-fold change (LFC ≥ |1|).

### Weighted gene co-expression network analysis (WGCNA)

WGCNA was performed on all samples except the *tlo*Δ/Δ using the *WGCNA* package (v 1.73) in R (Langfelder and Horvath 2008). Genes with fewer than fifteen counts in more than 75% of samples were excluded from the dataset, which left 3,963 of 6461 genes for further analysis. Genes (nodes) were assigned to co-expression modules, with a soft power threshold of 16, which sets the minimum R-squared value to 0.87 and the mean connectivity to 83.2. The remaining parameters used were as follows: TOMType = “signed”, mergeCutHeight = 0.1, numericLabels = FALSE, randomSeed = 1234, minModuleSize = 20, deepSplit = 3, verbose = 3. These parameters resulted in construction of 32 co-expression module eigengenes. *TLO* mutants were correlated to each of the 32 modules by a Pearson correlation analysis, with the correlation coefficients assigned p-values via Student’s t distribution test.

### Statistical Analysis

Statistical significance was calculated in R (v 4.4.1). For comparison of *TLO* mutants to the wildtype parental strain, the residuals values from an ANOVA test were first obtained. These values (degrees of freedom (df) and mean square (Mean sq)) were used in a Student’s t-test comparing each *TLO* mutant to the wildtype to correct for multiple tests. The standard error of the mean (SEM) and number of replicates were used to calculate the two-tailed p-value of each genotype compared to WT to assess significance. Genetic interaction among *TLO*s in the double mutant strains was determined via paired t-tests between predicted additive values (e.g. *tlo*α*9*Δ + *tlo*α*10*Δ) versus actual phenotype values (e.g. *tlo*α*9*Δ/*tlo*α*10*Δ) in each condition independently (Jafari and Ansari-Pour 2019; Mishra, et al. 2019).

### Data availability

The datasets generated during the current study are available from the corresponding author and 10.6084/m9.figshare.30890765. All WGS and RNA-Seq data are available from the NCBI Sequence Read Archive under Bioproject IDs PRJNA1305383 and PRJNA1117514.

## LIMITATIONS OF THIS STUDY

We acknowledge that this study does not mechanistically determine how phenotypes are produced by loss of single *TLO* genes. The explicit functions of each paralog remain unknown but are an active area of study. Phenotypes produced in mutants lacking a single *TLO* gene instead explore paralog redundancy and only provide circumstantial evidence for the role of individual paralogs in a specific cellular process. Secondly, we do not know how loss of single *TLO* genes alters the available pool of remaining Tlo proteins and their potential to be incorporated into Mediator. Thus, phenotypes could result from increased availability of other Tlo proteins that were not disrupted and are only an outcome of loss of a single *TLO* gene. However, it is important to note that the expression of a *TLO* gene was not predictive of phenotypic similarities across mutants, suggesting that the sequence identity of the disrupted *TLO* gene is also important. At the same time, the conclusions drawn from this study are influenced by the context of the investigation. The scope of investigated phenotypes and conditions for transcriptional profiling of the *TLO* mutants was not inclusive and our interpretations could change with greatly expanded phenotyping.

## ACKNOWLEDGEMENTS

We would like to thank Drs. Gary Moran and Derek Sullivan for multiple intellectual conversations regarding approaches to dissect *TLO* function. We would also like to thank members of the Anderson lab for fruitful discussions in designing this work and its format for presentation. Portions of R code written to generate figures and determine significance values were constructed with assistance from ChatGPT-4.

## AUTHOR CONTRIBUTIONS

Conceptualization: ES and MZA. Data curation: ES. Formal analysis: ES and NC. Funding acquisition: MZA. Investigation: ES, NC, MZ, PSH, ALW. Methodology: ES, ALW, and MZA. Project administration: MZA. Resources: MZA. Supervision: MZA. Validation: ES. Visualization: ES and MZA. Writing: ES and MZA.

