## Supplemental Figures for "Paralogs of the *Candida albicans TLO* gene family form interconnected functional networks with incomplete redundancy"

**SUPPLEMENTARY FIGURES**

**Fig. S1**

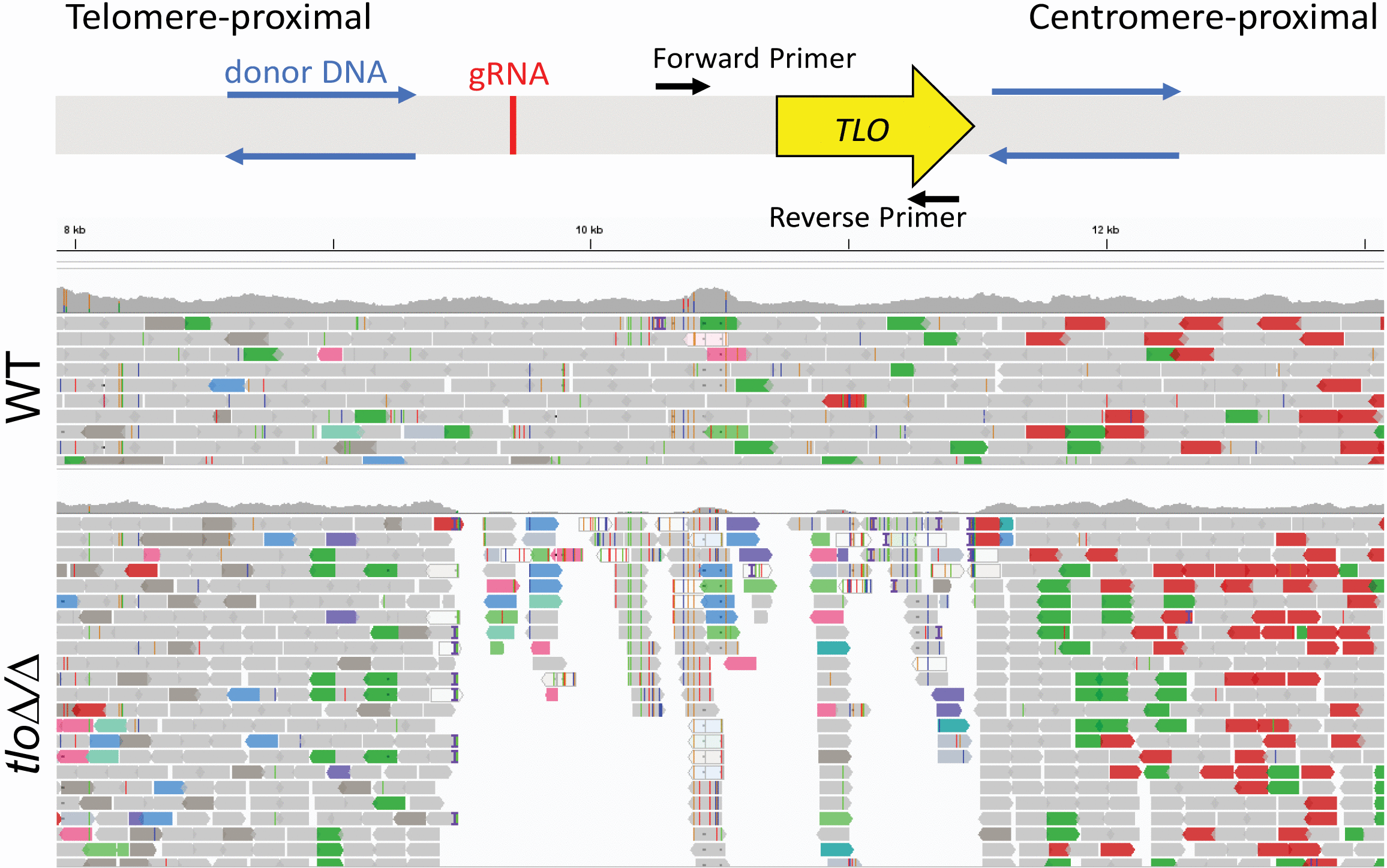

**Fig. S2**
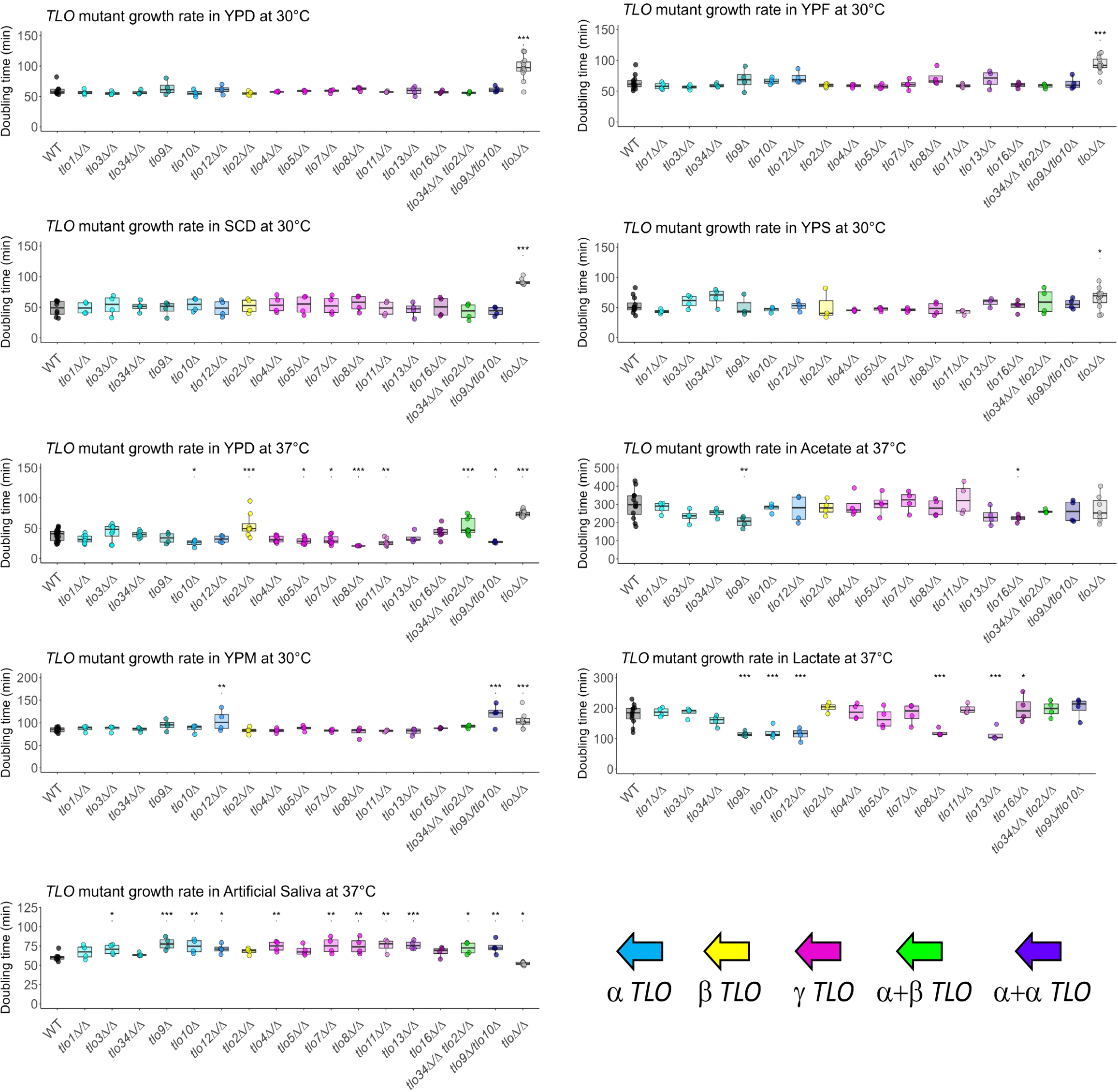

**Fig. S3**
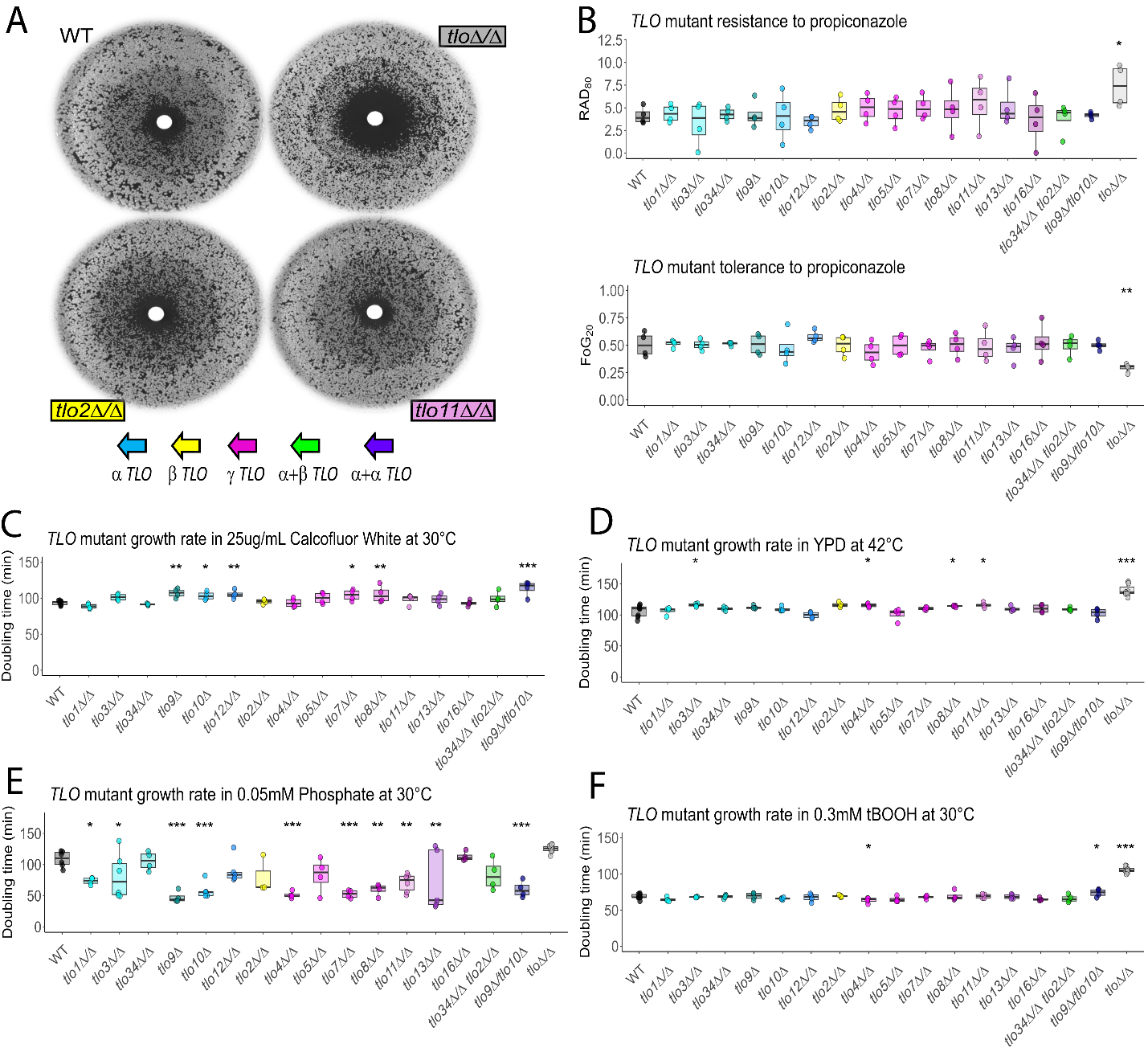

**Fig. S4**

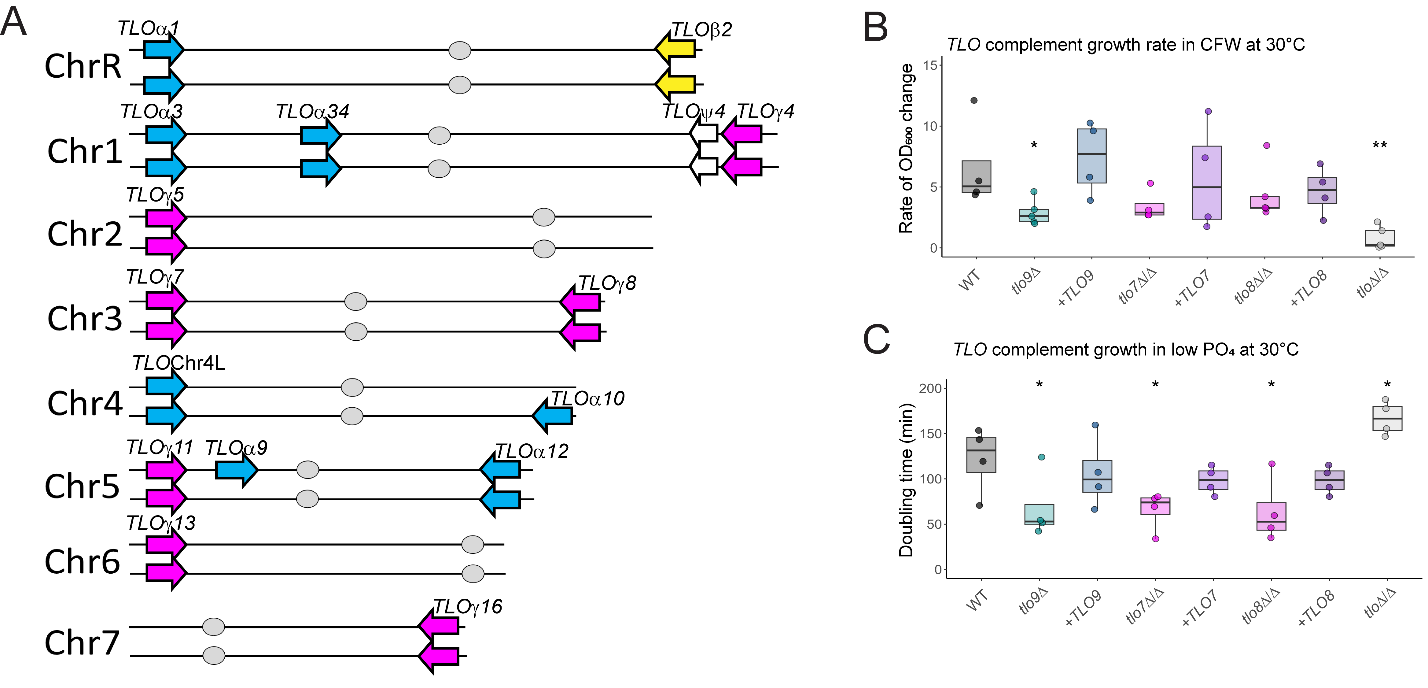

**Fig. S5**
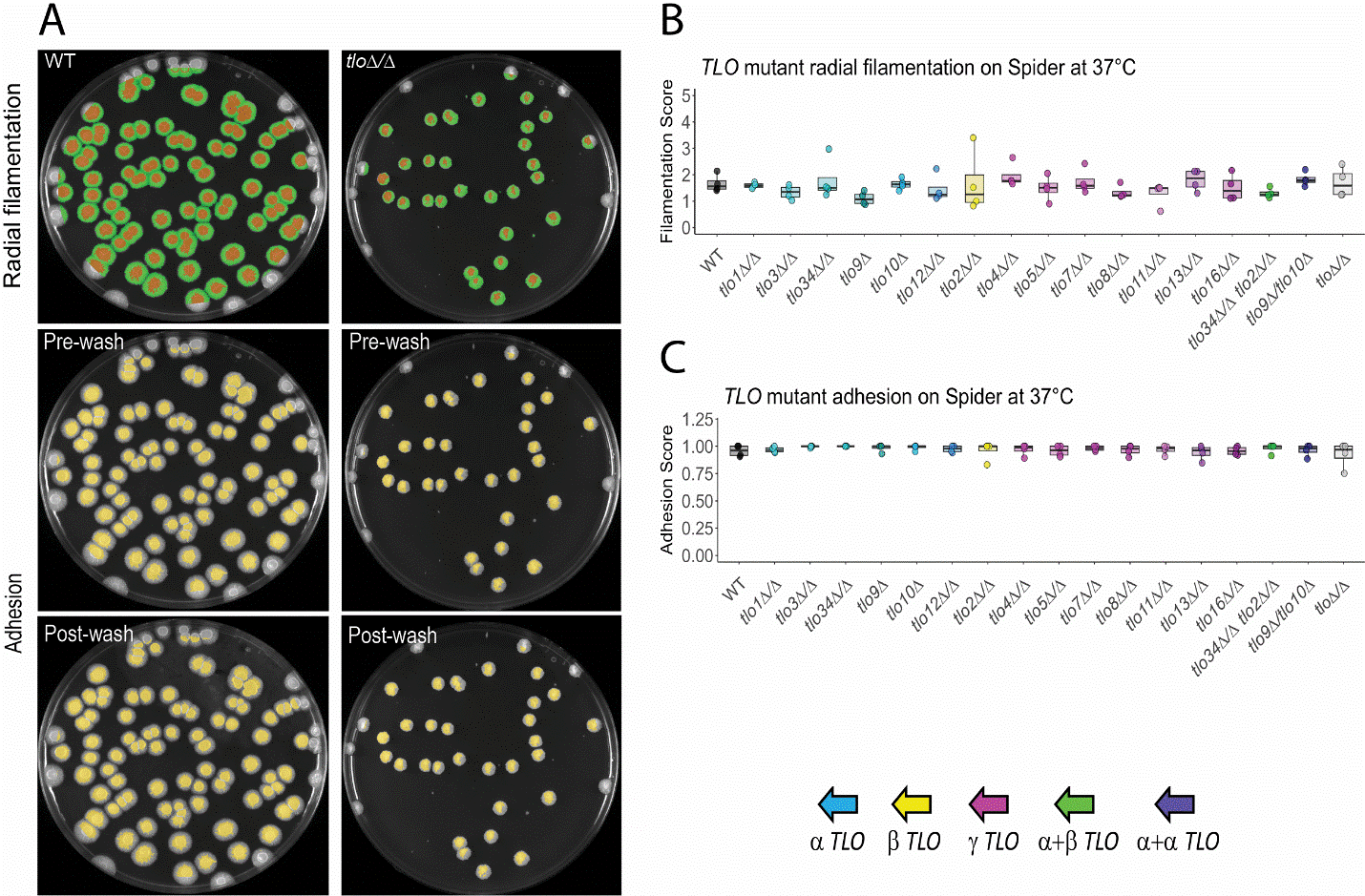

**Fig. S6**
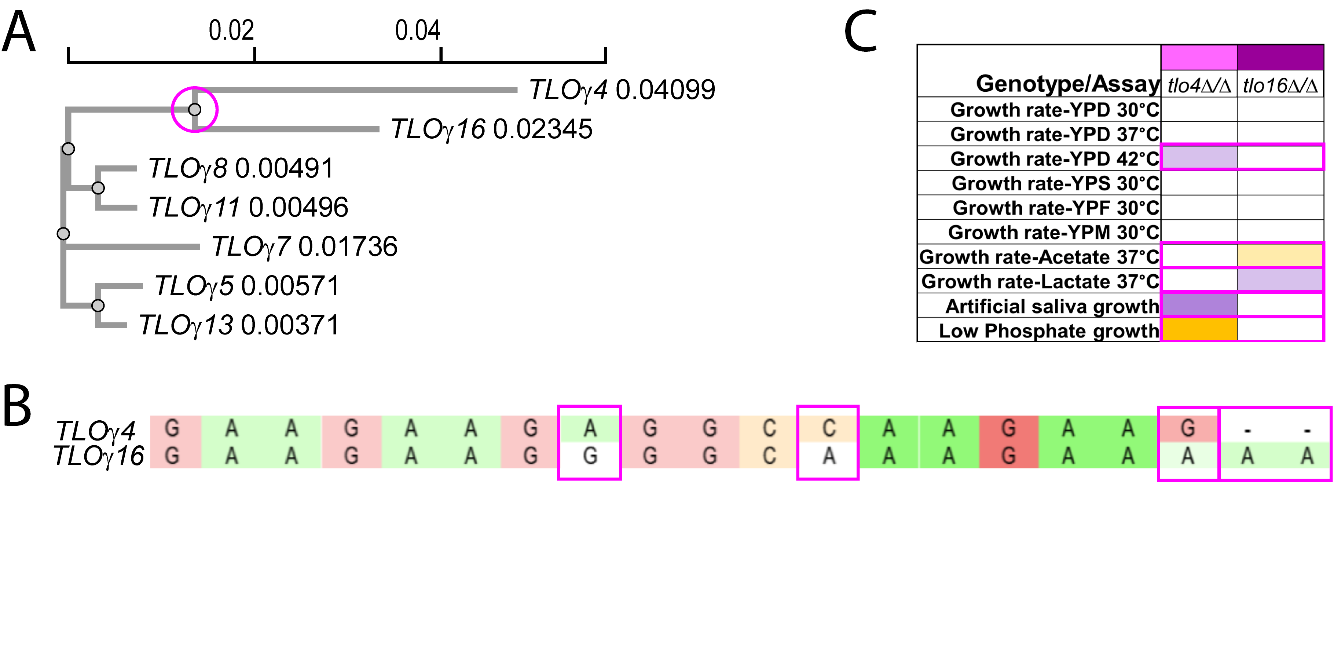

**Fig. S7**
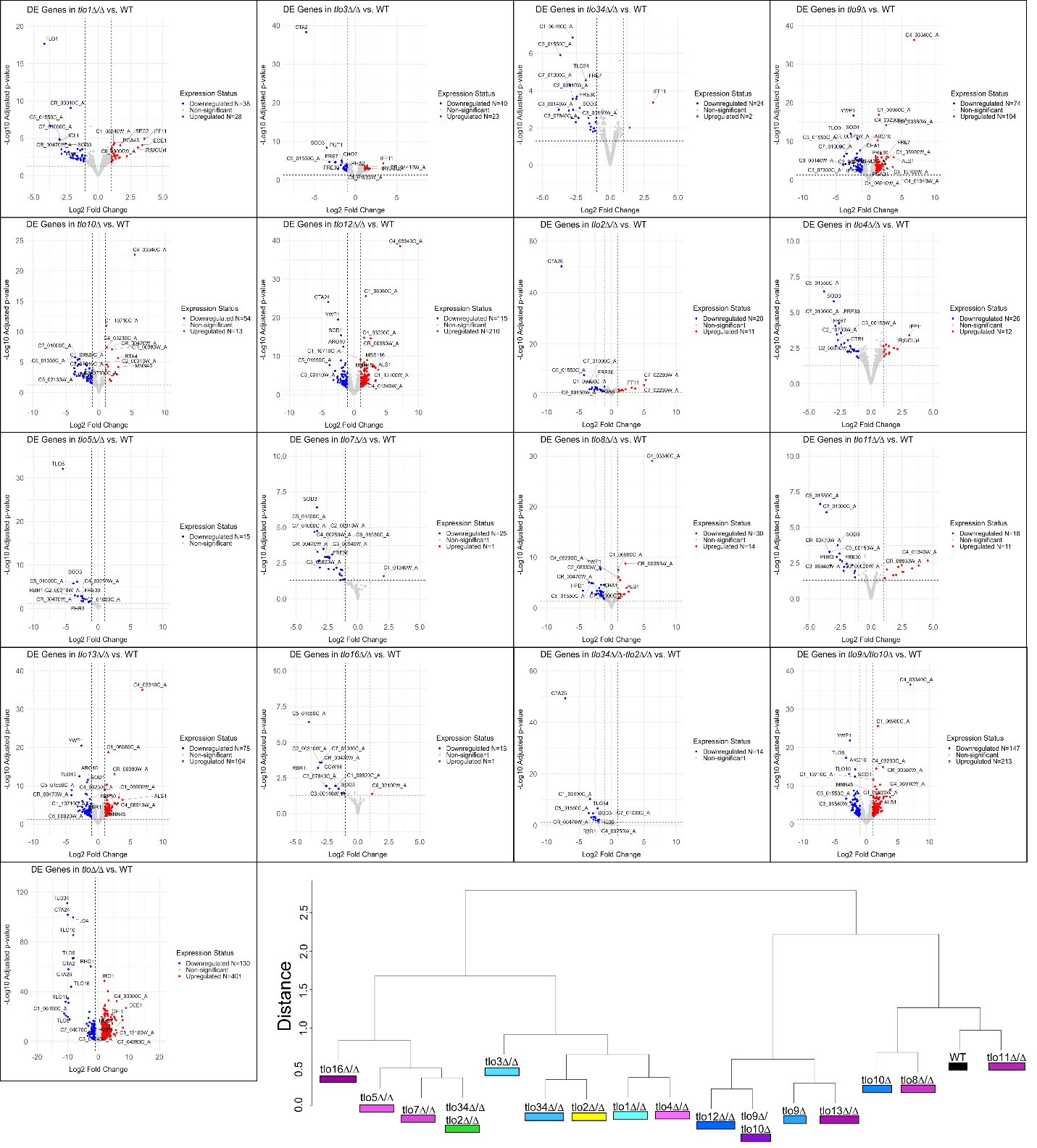

**Fig. S8**
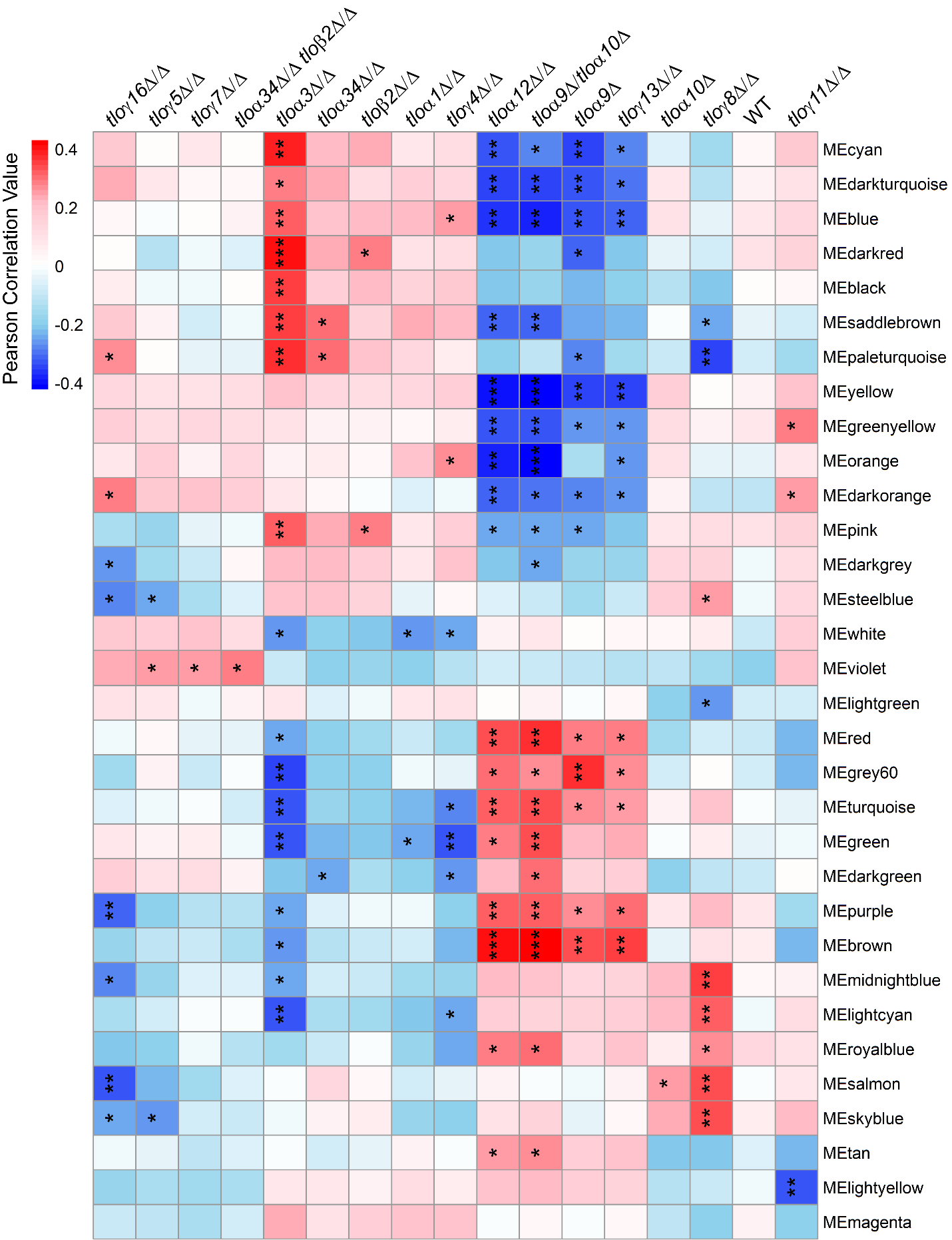

**Fig. S9**
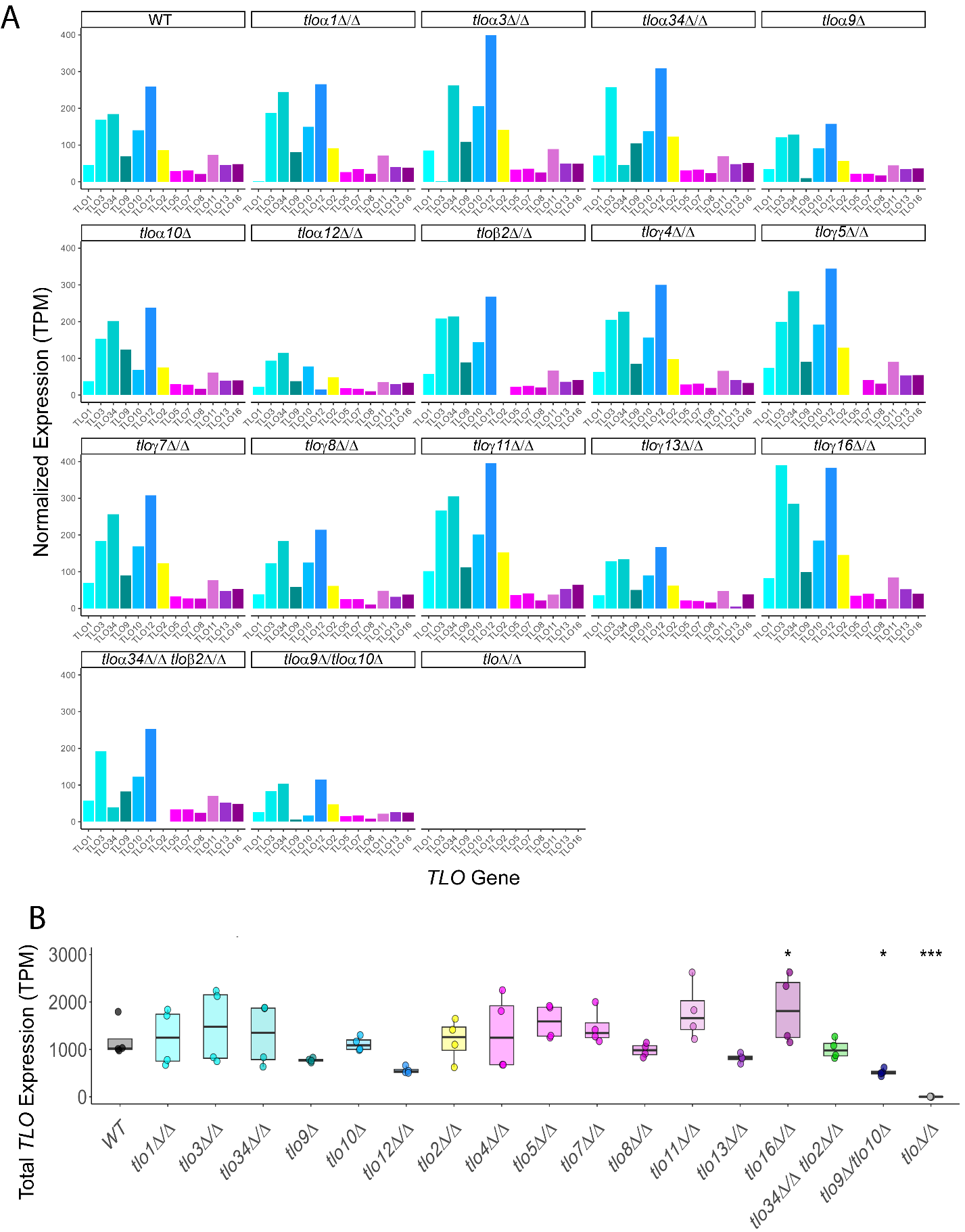

**SUPPLEMENTARY FIGURE LEGENDS**

**Fig. S1: Verification of constructed *TLO* mutants.** CRISPR-Cas9-mediated repair was used to delete individual *TLO* genes via a guide RNA targeted to unique surrounding sequence, with donor DNA of flanking regions used as a repair template. Two PCR checks were used to validate successful gene deletion with primers designed to target unique genomic DNA and *TLO*-specific 3’ sequences. Genomic DNA was extracted from putative knockouts, used to build gDNA libraries, then submitted for whole genome sequencing. With WT as the control, processed sequencing files were inspected manually in the Integrated Genome Viewer (IGV, v2.15.2) to verify on-target locus deletion as well as off-target effects such as long tract loss of heterozygosity.

**Fig. S1 alt text:** Graphical overview of the CRISPR/Cas9-mediated approach to construction of *TLO* gene deletions, with transcript coverage in the WT compared to a strain lacking a *TLO* targeted for deletion.

**Fig. S2: Growth of *TLO* mutants in various nutrient conditions.** With WT and *tlo*Δ/Δ strains as growth controls, all single and double *TLO* mutants were grown under an array of conditions, namely dextrose (YPD/SCD, artificial saliva), maltose (YPM), fructose (YPF), sucrose (YPS), acetate, and lactate, at either 30 or 37°C. *TLO* architectural groups are color-coded as indicated, and the *tlo*Δ/Δ strain is represented by light gray. A minimum of 4 biological replicates were used per condition. Significance was determined via modified Student’s t-tests; * p < 0.05, ** p < 0.01, *** p < 0.001.

**Fig. S2 alt text:** Growth data plotted as doubling time for all tested strains in nine different carbon sources and/or temperatures.

**Fig. S3: Analysis of mutant survival under environmental stressors.** (**A**) Cells were plated to YPD at an OD_600_ of 0.04 and allowed to dry for 1 hr. Sterile disks were inoculated with 25 μM of propiconazole and placed in the center of each plate. After incubation at 30°C for 48 hrs., images were captured. (**B**) Fraction of growth (FoG) and radius of inhibition (RAD) scores were calculated by the *diskImageR* (v1.1.0) package and each biological replicate was plotted. N=4. (**C-F**) Cells were grown under a variety of stressors (CFW, 42°C, low phosphate, and tBOOH) and doubling times were calculated from OD_600_ measurements by *growthcurver* (v0.3.1). N≥3. Significance was determined via modified Student’s t-tests; * p < 0.05, ** p < 0.01, *** p < 0.001.

**Fig. S3 alt text:** Graphics and data that show zones of clearance and associated resistance and tolerance scores for propiconazole in subfigures a and b. Growth data plotted as doubling times for all strains in four stress conditions, with subfigures labelled c to f.

**Fig. S4: Complementation of *TLO* mutant phenotypes.** (**A**) Single copies of *TLO*γ*7*, *TLO*γ*8*, and *TLO*α*9* were reintegrated into the respective *tlo*γ*7*Δ/Δ, *tlo*γ*8*Δ/Δ, and *tlo*α*9*Δ mutants at the *NEUT5L* locus. The *TLO*α*9* complement karyotype is displayed as a representative schematic. (**B**) Single *TLO* mutants and their complement strains were grown in CFW for 19 hrs. and (**C**) Low PO_4_ at 30°C for 30 hrs. N≥4. Significance was determined via modified Student’s t-tests; * p < 0.05, ** p < 0.01, *** p < 0.001.

**Fig. S4 alt text:** Graphics and data with an example karyotype of single TLO complementation at a neutral locus at chromosome 5 and subsequent doubling times or rate of OD change over time shown between WT, TLO mutants, and corresponding complement strains, with subfigures labelled a to c.

**Fig. S5: Measurement of solid filamentation phenotypes on Spider medium.** (**A**) Strains were plated for 100 CFUs onto solid Spider medium and grown at 37°C for 7 days. Images were captured before and after wash steps, then passed to the MIPAR (v5.1.0) software for quantification. (**B**) Radial filamentation scores were calculated with an equation ((area_hyphal growth_ – area_center colonies_) / (area_center colonies_)) based on pixel area reported by MIPAR. (**C**) Adhesion scores were calculated with the equation (area_colonies post-rinse_)/ (area_colonies pre-rinse_), in which the wash step entailed rinsing plates under a stream of water for 3-5 seconds in between imaging.

**Fig. S5 alt text:** Graphics and data from solid agar filamentation assays that demonstrate radial filamentation, adhesion, and invasion scores across strains, with subfigures labelled a to c.

**Fig. S6: Correlation between *TLO* sequence variation and mutant phenotypes.**

(**A**) A phylogenetic tree for the *TLO*γ architectural group was built by first aligning the nucleotide sequences with MUSCLE and then visualizing the sequence relationships using a distance-matrix phylogram. Relative evolutionary distance is marked on the phylogeny. (**B**) An alignment of the nucleotide sequences separated by the node marked in magenta using MUSCLE. Nucleotide differences are outlined in magenta. (**C**) Phenotypic differences for each single *TLO* mutant for the two paralogs compared in (B) are shown relative to the wildtype control. Phenotypic differences are outlined in magenta.

**Fig. S6 alt text:** Graphics representing the *TLO*γ phylogram and a subset of phenotype and nucleotide sequences between *TLO*γ*4* and *TLO*γ*16*.

**Fig. S7: Differentially expressed genes and transcriptional relatedness of *TLO* mutants.** DESeq2 was used to identify differentially expressed genes across all mutant strains versus WT, which were assembled into volcano plots. Significance and fold change thresholds are marked by dashed lines, with non-significant genes in light gray. Significantly up and down regulated genes are indicated with red and blue, with top DEGs for each genotype labelled. Significance was calculated by Wald test and corrected with Benjamini-Hochberg method; cutoffs: p < 0.05, LFC < |1|. Using WGCNA data, a dendrogram was used to cluster genotypes by closely related expression profiles.

**Fig. S7 alt text:** Graphics of differential gene expression data shown as volcano plots and a dendrogram of relative TLO relatedness based on gene expression data.

**Fig. S8: Co-expression driven analysis of *TLO* impact on cell functions.** (**A**) WGCNA was used to assemble co-expression module eigengenes (MEs) and correlate them to the mutant genotype. Significance calculated by Student’s t-distribution and corrected by the Benjamini-Hochberg method; * p < 0.05, ** p < 0.01, *** p < 0.001.

**Fig. S8 alt text:** A heatmap of Pearson’s correlation values between all strains except for the *tlo*Δ/Δ and gene co-expression modules built with WGCNA.

**Fig. S9: *TLO* expression across WT and mutant strains.** (**A**) Transcripts per million (TPM) values for each *TLO* were plotted for all *TLO* mutants. Bars are color-coded by group architecture., *TLO*γ*4* is excluded because it is misassigned in the Assembly 21 genome reference file. (**B**) The total TPM value for all *TLO* genes was calculated for each *TLO* mutant genotype and the wildtype. N=4. Significance was determined via modified Student’s t-tests; * p < 0.05, ** p < 0.01, *** p < 0.001.

**Fig. S9 alt text:** Graphics and data that show transcripts per million values for each *TLO* gene and total *TLO* expression across mutant and WT backgrounds, with subfigures labelled a and b.
